# [^18^F]DCP, First Generation PET Radiotracer for Diagnosis of Radiation Resistant Head and Neck Cancer

**DOI:** 10.1101/2020.04.28.063537

**Authors:** Xiaofei Chen, Kiran Kumar Solingapuram Sai, Zhe Li, Caigang Zhu, Kirtikar Shukla, Tom E. Forshaw, Hanzhi Wu, Stephen A. Vance, Megan Madonna, Mark W. Dewhirst, Allen W. Tsang, Leslie B. Poole, Nimmi Ramanujam, S. Bruce King, Cristina M. Furdui

## Abstract

Redox metabolism plays essential functions in the pathology of cancer. As tumor redox profiles uniquely reflect cancer stage and in select cases, therapeutic sensitivity, the capability to image redox molecular features is essential to improve diagnosis, treatment, and overall quality-of-life (QOL) of cancer patients. While a number of radiotracers for imaging redox metabolism have been developed, there are no reports of radiotracers for in vivo imaging of protein oxidation. Here we take the first step towards this goal and describe the synthesis and kinetic properties of a new positron emission tomography (PET) [^18^F]DCP radiotracer for in vivo imaging of protein sulfenylation. Time course biodistribution and PET/CT studies using xenograft animal models of Head and Neck Squamous Cell Cancer (HNSCC) demonstrate feasibility of diagnosing radiation resistant tumors, which display lower [^18^F]DCP signal. These findings are consistent with our previous reports of decreased protein sulfenylation in clinical specimens of radiation resistant HNSCC. We anticipate further development and implementation of this concept in clinical practice to improve the diagnosis of patients with radiation resistant tumors and the accuracy of prognosis for patients undergoing radiation treatment.

**Single Sentence Summary:** The study introduces a new PET radiotracer for profiling tumor protein oxidation as a prognostic indicator of resistance to radiation therapy.

## Introduction

Approximately 3.5 million cancer patients worldwide are treated with radiation therapy every year. These include Head and Neck Squamous Cell Cancer (HNSCC) patients, where radiation alone or combined with systemic chemotherapy is applied either as definitive or as adjuvant post-surgical therapy. Toxicity associated with these harsh treatments has significant consequences on breathing, swallowing, and speech functions affecting long-term quality of life. Even when treatment is successful, HNSCC patients often suffer sustained life-long sequelae such as xerostomia, dysphagia, dependence on a gastric tube for nutrition, neck muscle spasms and contractions with significant neck pain, tracheostomy tube, dysphonia, and others. As a result, the suicide rates in HNSCC patients are among the highest (50.5/100,000 person-years) in cancer patients, second only to lung cancer patients (81.4/100,000 person-years) (average rate in US population is 12/100,000 person-years) (1).

Thus, there is an immediate need for strategies to reduce acute and long-term toxic effects of radiation therapy on healthy tissue, and to identify patients with radiation resistant tumors early on so they can be spared from unproductive radiation therapy. These patients could undergo treatment with other potentially life-saving therapies before valuable time is irreversibly lost. As one of the mechanisms by which radiation exerts its effects is through the generation of reactive oxygen species (ROS) (2), redox metabolism has become central to the quest for the discovery of new solutions to these challenges. Indeed, redox-active compounds such as Fe- and Mn-porphyrins (3–5), small molecule thiol precursors (e.g., Ethyol (amifostine)) (6), and alternative anti-inflammatory therapies (7) have been shown to protect healthy tissue from radiation-induced damage and are utilized to alleviate oral mucositis and xerostomia in HNSCC patients. Identification of patients with radiation resistant tumors has been, however, more challenging. While differential [^18^F]FDG positron emission tomography (PET) scans collected before and after radiation treatment could predict the risk of tumor recurrence (8, 9), this approach still requires patients to undergo first treatment with ionizing radiation.

Redox metabolism has been known for many decades as a key mediator of the response to radiation treatment with radiation resistant cells and tumors being characterized, in general, by a pattern of low basal ROS and upregulated antioxidant systems, consistent with our findings in HNSCC (10, 11). One of the well-established mechanisms of ROS signaling is through oxidation of a select class of protein cysteine thiols resulting in stable or transient sulfenylation of proteins (-SOH) (12). Intra- or intermolecular protein disulfides (-SS-), persulfides (-S_n_H), sulfenamides (-SN), sulfinic and sulfonic acids (-SO_2_H and -SO_3_H, respectively) and other species are formed upon the reaction of sulfenylated species with other cysteines, H_2_S, or excess ROS (13). Due to their reversibility, these oxidative modifications are ideal protein molecular switches implicated as central mechanisms in the cellular sensing of redox microenvironment, and as regulators of a multitude of processes vital to cellular survival and proliferation.

To facilitate the investigation of these redox processes, we reported on synthesis and applications of a series of chemical probes for selective labeling of sulfenylated proteins under both *in vitro* and *in vivo* conditions over the past years (14–18). These compounds include BP1, a biotin-tagged derivative of (2,4-dioxocyclohexyl)propyl (DCP) (17), which was crucial to our discovery of suppressed protein sulfenylation in clinical specimens of radiation resistant HNSCC (10, 11). This finding formed the basis of the strategy presented here, where we envisioned that selective profiling of protein sulfenylation could be applied to identify patients most likely to benefit from radiation therapy and to identify cases where redox-altering adjuvants could be combined with radiation therapy to improve clinical outcomes.

Towards this goal, we report the synthesis of the first positron emission tomography (PET) radiotracer [^18^F]DCP for detection of sulfenylated proteins and its successful application to differentiate radiation resistant from radiation sensitive tumors in xenograft animal models of HNSCC. The proof-of-concept studies were performed using a matched model of radiation response for HNSCC, radiation sensitive SCC-61, and radiation resistant rSCC-61 mimicking radiation response profiles in recurrent tumors. *In vivo* biodistribution and PET imaging measurements were then extended to include HNSCC tumors generated from radiation sensitive JHU022 and radiation resistant SQ20B cell lines. The overall findings validate the capability of [^18^F]DCP to phenotype radiation sensitivity of HNSCC tumors.

## Results

### Synthesis and kinetic characterization of [^18^F]DCP and non-radioactive [^19^F]DCP

Given the short half-life of ^18^F (t_1/2_ 109.7 min) and the projected application of [^18^F]DCP in clinical imaging, the synthesis of this derivative needed to meet a number of requirements: follow a simple and quick synthesis scheme, proceed with high yield, and be compatible with procedures, reagents and equipment commonly available at hospital PET centers. Following these needs, we synthesized both the radioactive [^18^F] and non-radioactive [^19^F] versions of DCP in 2-3 reaction steps (**Figure 1, A and B**). Detailed synthetic procedures and characterization of chemical species can be found in the Materials and Methods and Supplementary Material, and are briefly summarized next. First, the copper-catalyzed click reaction of the previously described DCP-alkyne (19) with tosylated azide, prepared in two steps from 2-bromoethanol (20) (Supplemental **Figure S1, A and B**), produced the protected tosylated triazole (**1**, **Figure 1A**; Supplemental, **Figure S2, and B**) in 74% yield. Ceric ammonium nitrate oxidation of **1** further generated the requisite precursor **2** for [^18^F] incorporation in 73% yield (Supplementary Material, **Figure S3A and B**). From here, the automation of [^18^F]DCP synthesis was carried out on a TRASIS AIO radiochemistry module (n=10) at the Wake Forest PET Research Facility yielding a final compound of high radiochemical purity (>98%), high specific activity ~2,610-2,820 mCi/μmol, and a radiochemical yield of 18% (decay corrected to end of synthesis). Radiochemical purity of the tracer was verified by co-injection of the non-radioactive standard [^19^F]DCP and HPLC analysis (Supplemental **Figure S4**). Radiochemical synthesis, including [^18^F]^−^ reaction, HPLC purification, and radiotracer formulation was completed within 40-45 min. The cold [^19^F]DCP was similarly prepared starting with DCP-alkyne and copper-catalyzed click reaction with the previously described fluorinated azide (21) to generate first a protected fluorinated triazole (**3**, **Figure 1B**; Supplemental **Figure S5, A and B**) in 71% yield. This was followed by ceric ammonium nitrate deprotection to generate [^19^F]DCP in 70% yield (**Figure 1B**; Supplemental **Figure S6, A and C**).

**Figure 1.**
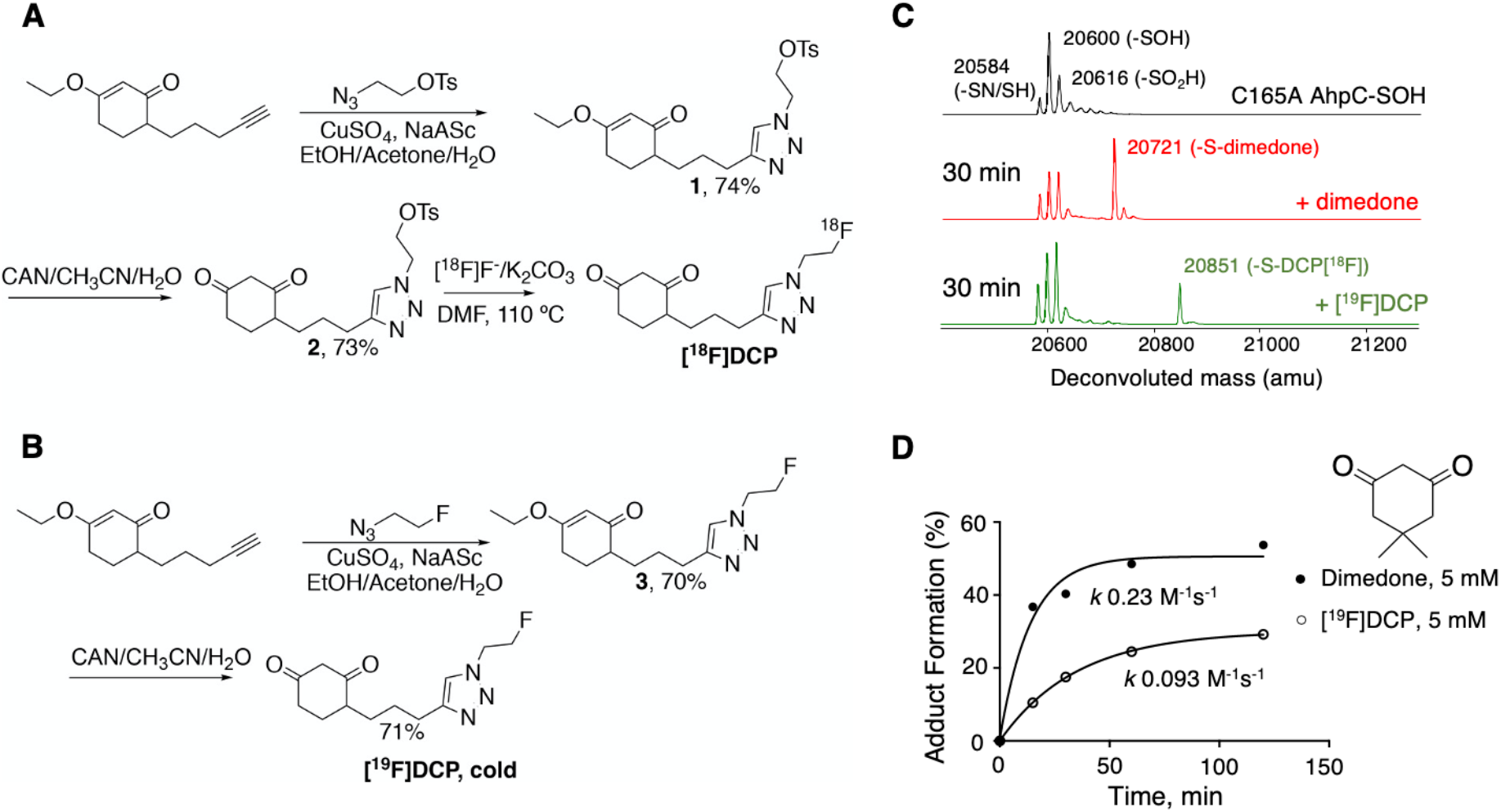
Synthesis and kinetic characterization of 1^st^ generation protein oxidation tracer [^18^F]DCP. **(A)** Synthesis scheme for [^18^F]DCP. **(B)** Synthesis scheme for cold [^19^F]DCP. **(C)** Deconvoluted mass spectra of oxidized C165A AhpC (black), and its 30 min reaction with 5 mM dimedone (red) or [^19^F]DCP (green). **(D)** Kinetic profiles and reaction rates of oxidized C165A AhpC reaction with 5 mM dimedone (black circle) or [^19^F]DCP (open circle). The data were fit in GraphPad Prism 7.0 using single exponential kinetics and the second order rate constants were calculated by dividing the rate with the respective concentration of probes (5 mM).

Next, we investigated the selectivity and kinetics of the new [^19^F]DCP derivative towards sulfenylated proteins, using a model recombinant protein, C165A AhpC, and procedures employed previously for development of other protein sulfenylation probes (2, 14–18). C165A AhpC is a mutant of *Salmonella typhimurium* peroxiredoxin AhpC, which stabilizes mixed reduced (-SH) and oxidized sulfenic (-SOH), sulfenylamide (-SN) and sulfinic (-SO_2_H) species at the reactive cysteine C46 enabling evaluation of [^19^F]DCP reactivity with oxidized proteins. The results in **Figure 1C** show [^19^F]DCP retains selectivity for sulfenylated C165A AhpC, and displays ~2-fold lower kinetics relative to dimedone, the closest structural analog of the DCP core in [^19^F]DCP (dimedone: *k* 0.23 M^−1^s^−1^; [^19^F]DCP: *k* 0.093 M^−1^s^−1^; **Figure 1D**).

### [^18^F]DCP serum stability and in vitro cellular uptake

Serum stability is a key qualifier for any diagnostic or therapeutic agent. The serum stability of [^18^F]DCP was assessed by incubation with human serum and HPLC analysis of aliquots extracted at different time points. The data indicate minimum degradation over a 4 h time course with ~95% of [^18^F]DCP remaining intact at the 4h time point (**Figure 2A**). Having demonstrated the compatibility with *in vivo* applications from the perspective of serum stability, the next studies focused on the cellular binding efficiency and selectivity of [^18^F]DCP using the matched SCC-61 and rSCC-61 HNSCC cells, which have been thoroughly characterized and reported in previous publications (10, 11, 22, 23). Compared with the radiation sensitive SCC-61, the radiation resistant rSCC-61 cells display increased resistance to ionizing radiation (SCC-61: SF2 0.35, rSCC-61 SF2 0.66; SF2 - survival factor at 2 Gy), decreased ROS including cytosolic, mitochondrial and nuclear H_2_O_2_, upregulation of antioxidant proteins (e.g., Prx 1-3, GST-pi, NQO1) (10), increased mitochondrial MTHFD2 contributing to higher NADPH levels (23), and increased glucose uptake with glucose metabolism being funneled into the Pentose Phosphate Pathway and leading overall to increased NADPH/NADP^+^ ratio in these cells (SCC-61: 11.1; rSCC-61: 17.5) (11). The rSCC-61 cells were also characterized by decreased mitochondrial respiration measured by the oxygen consumption rates (SCC-61: 169 pmoles/min; rSCC-61: 96 pmoles/min) and ATP synthesis (SCC-61: 155 pmoles/min; rSCC-61: 83 pmoles/min) (11).

**Figure 2.**
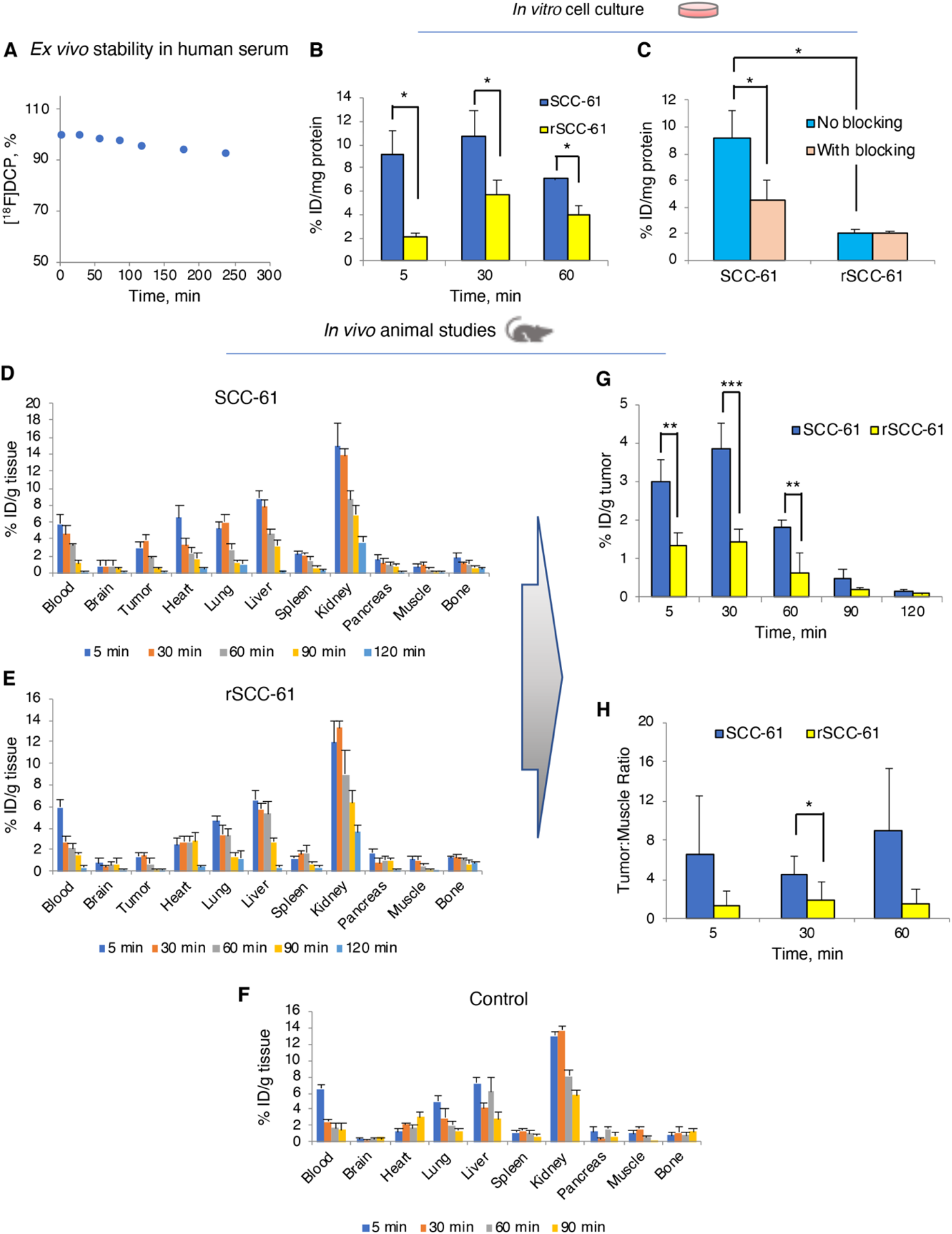
Primary characterization of [^18^F]DCP compatibility with *in vivo* applications. (**A**) *Ex vivo* serum stability assay for [^18^F]DCP performed by incubating the radiotracer with human serum and quantifying by HPLC analysis over a 4h time course. (**B**) *In vitro* cell uptake of [^18^F]DCP in SCC-61 and rSCC-61 cells after 5 min, 30 min and 60 min of radiotracer exposure. (**C**) *In vitro* assessment of binding specificity using blocking with non-radioactive [^19^F]DCP analog in SCC-61 and rSCC-61 cells. Blocking of sulfenylated proteins was demonstrated by exposing the cells to 50x excess non-radioactive [^19^F]DCP, 15 min prior to adding the radiotracer. (**D**) **-**(**H**) Biodistribution of [^18^F]DCP in matched xenograft animal models of radiation response: radiation sensitive SCC-61 (**D**), radiation resistant rSCC-61 (**E**), and control non-tumor carrying animals (**F**). (**G**) Analysis of biodistribution data shows accumulation of [^18^F]DCP in SCC-61 tumors 2-3 fold more than in rSCC-61 tumors, a ratio stable up to 90 min post injection. (**H**) Tumor to muscle ratios extracted from the biodistribution data show [^18^F]DCP accumulation in SCC-61 tumors compared to rSCC-61 tumors. The data in (B) - (G) were expressed as % injected dose (ID)/mg of protein present in each well with p values ≤ 0.05 considered statistically significant (n=3). Statistical analysis (t-test) was performed using Microsoft Excel and all results are displayed as mean ± standard deviation. Asterisks indicate statistically significant changes [α = 0.05, p values of 0.01–0.05 (*), 0.001– 0.01 (**), or <0.001 (***)].

Given the richness of information available for this system matching general molecular features that distinguish radiation resistant cells and tumors across HNSCC, the SCC-61 and rSCC-61 cells were utilized in the initial studies to evaluate the capability of [^18^F]DCP to preferentially label radiation sensitive cells and tumors. The first studies measured the uptake of [^18^F]DCP in SCC-61 and rSCC-61 cells using the non-radioactive [^19^F]DCP as a blocking agent. The [^18^F]DCP signal at 5, 30 and 60 min incubation times was 5-, 2.5-, and 2-fold higher, respectively, in SCC-61 compared to the rSCC-61 cells (**Figure 2B**), consistent with the higher ROS and protein sulfenylation in SCC-61 cells (10). Pre-treatment of SCC-61 cells for 15 min with non-radioactive [^19^F]DCP, decreased the radioactive signal by ~60%, while no significant blocking of [^18^F]DCP was observed in the rSCC-61 cells (**Figure 2C**), presumably due to the already low [^18^F]DCP signal in these cells.

### In vivo evaluation of [^18^F]DCP using SCC-61 and rSCC-61 derived tumor xenografts

Time-course biodistribution studies using SCC-61 and rSCC-61 xenograft animal models were performed to validate *in vivo* the findings from the cell culture studies. Mice were sacrificed at 5, 30, 60, 90 and 120 min (n = 4 for each time point) after the injection of [^18^F]DCP. Multiple organs (brain, heart, lung, liver, spleen, pancreas, and kidney), tissues (muscle, tumor), bone and blood were removed, weighed, and residual radioactivity was counted using an automatic gamma-counter. The radiotracer uptake was calculated as percentage of injected dose per gram of biological specimen (tumor, organ, tissue, bone and blood; %ID/g) (**Figure 2, D-F**). The radioactivity peaked at 30 min in the tumor and gradually washed out from the blood by 120 min. Kidney being the primary excretory organ had the highest residual uptake at 120 min. Importantly, there was no significant bone uptake, suggesting lack of defluorination. Focusing on the tumor, the data showed >60% higher [^18^F]DCP in the radiation sensitive SCC-61 relative to the radiation resistant rSCC-61 tumors (**Figure 2G**). The results also showed good tumor-to-muscle ratio with >80% enrichment in the tumor (**Figure 2H**) encouraging further investigations.

### Metabolic and vascular characteristics of the SCC-61 and rSCC-61 xenograft tumors

Additional studies were performed to verify that the *in vitro* redox profiles of SCC-61 and rSCC-61 cells were maintained *in vivo* and to ensure that the differential labeling of the radiation resistant and sensitive tumors by [^18^F]DCP was not due to differences in tumor perfusion. As noted above, the rSCC-61 cells display higher glucose uptake and lower mitochondrial activity compared with SCC-61 cells (11). Both glucose uptake and mitochondria membrane polarization in SCC-61 and rSCC-61 tumors were characterized using quantitative optical spectroscopy (24, 25). Glucose uptake was monitored with 2-NBDG (2-(*N*-(7-nitrobenz-2-oxa-1,3-diazol-4-yl)amino)-2-deoxyglucose, a fluorescent analog of glucose) and mitochondria membrane polarization with TMRE (tetramethylrhodamine ethyl ester, a cationic fluorescent dye binding to active mitochondria). Importantly, to avoid artifacts associated with analysis of tumors of different volumes, SCC-61 and rSCC-61 tumors with a similar diameter of ~6 mm were selected for analysis. The results in **Figures 3** show that there was indeed higher 2-NBDG uptake and lower TMRE staining in the rSCC-61 than in the SCC-61 tumors consistent with the previous in vitro findings of increased glucose uptake and decreased mitochondrial respiration in rSCC-61 cells (11). Furthermore, the analysis of baseline oxygen saturation and hemoglobin [Hb] levels show these to be comparable between the two tumors. Cumulatively, these results demonstrate that noted differences in [^18^F]DCP incorporation are not due to lower perfusion or hypoxia of rSCC-61 tumors but to differences in protein sulfenylation.

**Figure 3.**
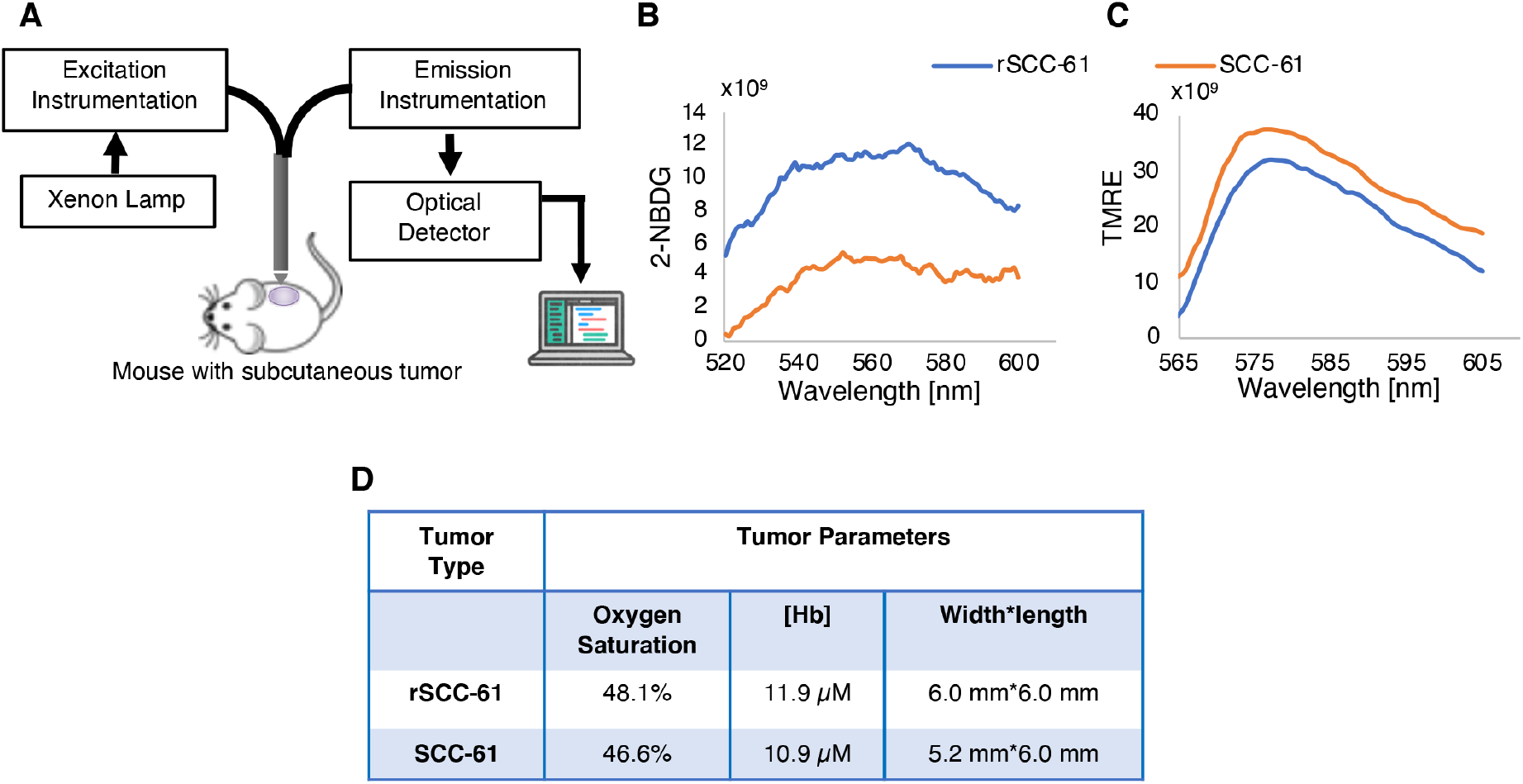
Quantitative optical spectroscopy tumor measurements. (**A**) Key elements of experimental setup. (**B**) Glucose uptake: Quantification of 2-NBDG fluorescence signal showing higher uptake in rSCC-61 compared with SCC-61 tumors. (**C**) Mitochondria membrane polarization: Quantification of TMRE fluorescence signal shows lower labeling of rSCC-61 compared with SCC-61 tumors. (**D**) Quantification of baseline oxygen saturation and [Hb] shows comparable levels between the rSCC-61 and SCC-61 tumors.

### MicroPET/CT imaging and biodistribution in a panel of HNSCC tumors

To further demonstrate *in vivo* the ability of [^18^F]DCP to discriminate between radiation resistant and sensitive tumors, the biodistribution studies were extended to include two additional HNSCC cell lines, the radiation sensitive JHU022 (SF2 0.2) and radiation resistant SQ20B (SF2 0.77). Thus, while the matched SCC-61 and rSCC-61 cell lines mimic recurrence of radiation resistant tumors in the same patient, and SQ20B and JHU022 represent radiation resistant and sensitive cells, respectively, from genetically unique patients. For these studies, we focused specifically on Human papillomavirus (HPV) negative HNSCC to match the clinical need given the increased prevalence of vaping worldwide (26, 27) and the known increased sensitivity to radiation therapy of HPV positive HNSCC (28, 29).

Subcutaneous tumors were generated in mice by injecting the cells between the shoulder blades to avoid potential interference from the bladder during microPET imaging and using procedures described in Materials and Methods. While orthotopic oral implantation of cells for these studies would have been preferable, this was ultimately not feasible due to concerns regarding the animals’ welfare and experimental limitations (e.g., the tumors can only grow a few millimeters before the animals cannot eat, ease of access for rapid tumor dissection for the time course biodistribution studies, and other considerations). After the presence of the tumor was confirmed through bioluminescence (**Figure 4A**, top images), [^18^F]DCP was injected through the tail vein and the tumors were imaged using microPET at 50 min post-injection (100 ± 20 μCi of [^18^F]DCP, 20 min scan time, n = 6). Representative axial and sagittal microPET/CT images are shown in **Figure 4A** middle and bottom panels, respectively. Data analysis showed a trend where the radiation sensitive tumor had increased average ROIs relative to the radiation resistant tumors: SCC-61 > JHU022 > SQ20B > rSCC-61.

**Figure 4.**
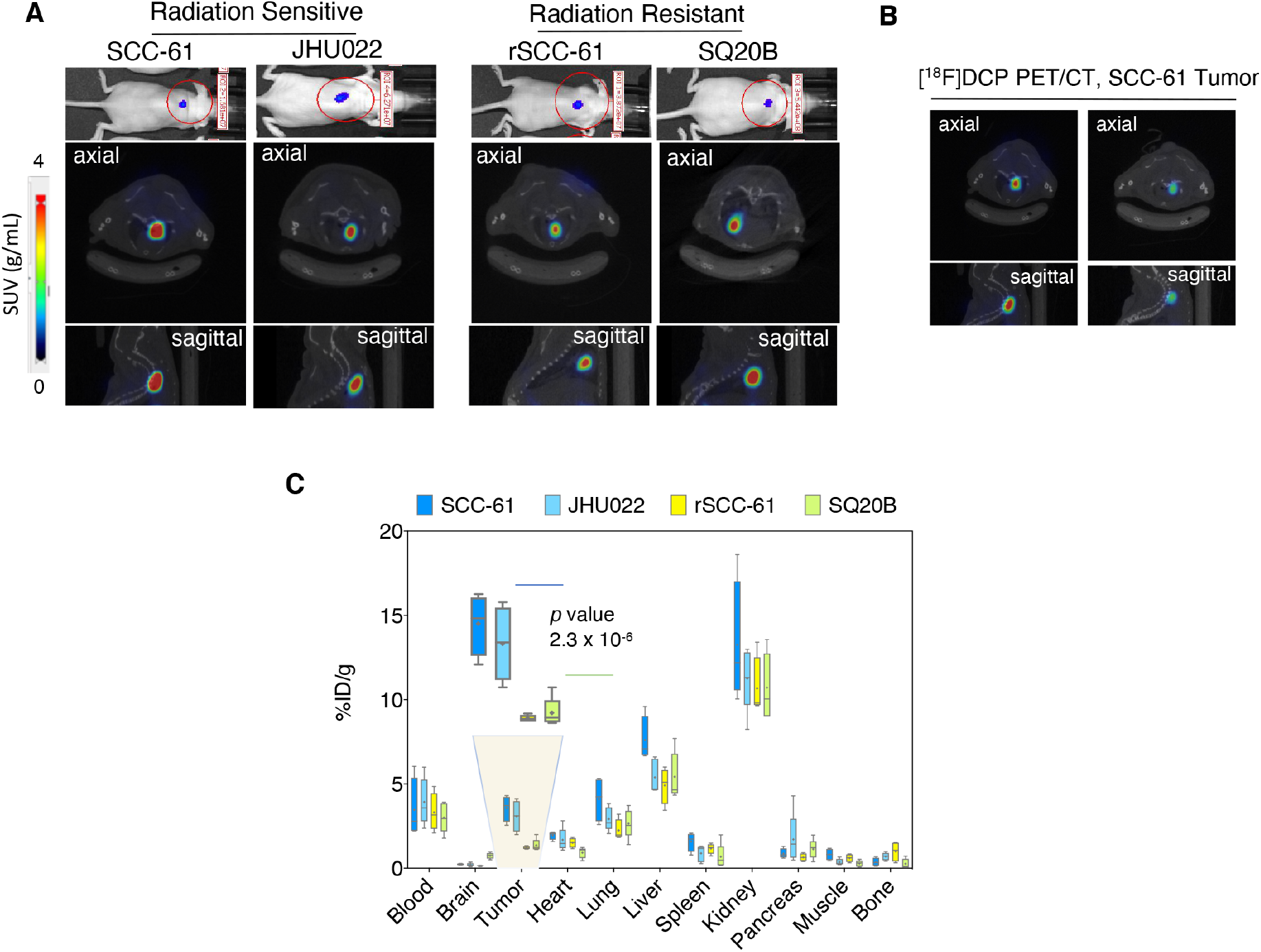
*In vivo* microPET/CT and biodistribution studies with [^18^F]DCP. (**A**) Representative IVIS and coronal fused microPET/CT images of [^18^F]DCP in HNSCC tumor bearing mice at 50 min post injection. The corresponding IVIS luciferase imaging is shown at the top. MicroPET ROI analysis shows higher signal intensity in the radiation sensitive SCC-61 and JHU022 tumors compared with the radiation resistant rSCC-61 and SQ20B tumors. SUV: standardized uptake value. (**B**) Blocking of radiotracer signal is demonstrated using SCC-61 tumors by injecting the cold ligand 20 min prior to the injection of the radiotracer. (**C**) Biodistribution analysis shows [^18^F]DCP accumulates in the radiation sensitive SCC-61 and JHU022 tumors 2.5 fold more than in rSCC-61 and SQ20B tumors at 60 min post injection. Statistical analysis (t-test) was performed using Microsoft Excel and all results are displayed as mean ± standard deviation.

Two SCC-61 tumor-bearing mice from the same cohort were used for blocking experiments with cold [^19^F]DCP (**Figure 4B**). These two mice received [^19^F]DCP, 10 mg/kg, 20 min prior to the injection of [^18^F]DCP. The axial and sagittal microPET/CT images shown in **Figure 4B** demonstrate significant blocking of the radiotracer signal by pre-treatment with cold [^19^F]DCP, evidencing *in vivo* the high selectivity of the radioligand. Notably, there was no toxicity noted with the injection of cold [^19^F]DCP at 10 mg/kg, a dose 36,000-fold higher than the dose of [^18^F]DCP corresponding to 0.27 μg/kg in these blocking studies.

To more precisely quantify the level of radioactivity in the tumors and across the mouse anatomy, we conducted biodistribution studies similar to those shown in **Figure 2**, with the difference that the analysis was performed at a single time point, 60 min post-injection of [^18^F]DCP (**Figure 4C**). Overall, the biodistribution analysis showed a statistically significant 2.5 fold increased [^18^F]DCP in the radiation sensitive SCC-61 and JHU022 tumors compared with the radiation resistant SQ20B and rSCC-61 (*p* 2.3 × 10^−6^). In general, the larger radioactive signal in the kidney is expected with 60 min post-injection radiotracer analysis.

## Discussion

Initially regarded as harmful by-products of cellular aerobic metabolism, ROS have been increasingly appreciated as pivotal signaling molecules that regulate a spectrum of physiological and pathological processes. It is important that cells carefully balance the production and the elimination of ROS to maintain a physiologically healthy redox environment (30). Elevated ROS have been observed in nearly all cancers and are thought to promote tumor initiation and progression by mediating genome instability, cell survival and proliferation, angiogenesis, and metastasis (31–34). This shift to a more oxidative state may render cancer cells vulnerable to ROS manipulation and many cancer therapies exploit this vulnerability by either increasing ROS generation or decreasing ROS scavenging (33–35).

Effective treatment of cancer with ionizing radiation also relies on accumulation of ROS and other species leading to DNA damage and cell death (36, 37), in most cases with the exception of tumors harboring enhanced antioxidant capacity, which are resistant to treatment (38–42). The new PET imaging approach presented here takes advantage of these unique redox metabolic features of radiation sensitive and resistant tumors to stratify patients based on their likelihood of response before the start of treatment. While several PET imaging probes were developed over the years that either react with ROS directly (e.g., ascorbic acid (43, 44), and our recent work (45)) or are activated by redox chemistry (e.g., hypoxia imaging agents ATSM, FMISO and FAZA (46)), these have limited value as predictors of radiation response *in vivo* as the relationship between oxygen saturation and ROS is quite complex and low oxygen saturation does not necessarily mean low ROS or protein oxidation as also exemplified by the spectroscopy imaging data presented here.

Our published studies using a matched model of radiation response for HNSCC found significantly decreased levels of ROS and increased levels of antioxidant proteins in the radiation resistant rSCC-61 cells compared with the radiation sensitive SCC-61 (10, 11). The enzymatic activity of antioxidant proteins was supported in rSCC-61 cells by increased production of NADPH, which is critical to sustain the cycling of cellular peroxiredoxins, major antioxidant enzymes upregulated in radiation resistant cells and involved in degradation of H_2_O_2_ and lipid peroxides (23). Overall, differences in redox metabolism are reflected in the DNA damage and protein oxidation profiles, with the radiation resistant rSCC-61 cells displaying lower level of single and double-strand DNA breaks and lower protein sulfenylation, a finding validated using *ex vivo* analysis of human HNSCC tumor specimens (10).

Analysis of protein sulfenylation in these studies was enabled by development of (2,4-dioxocyclohexyl)propyl (DCP) family of sulfenic acid-selective chemical probes (47), which now include a variety of derivatives and tags such as: fluorophores (DCP-FL1, DCP-FL2) (16), biotin (DCP-Bio1, DCP-Bio2, DCP-Bio3 (16); BP1, containing the reactive 1,3-dicarbonyl DCP component (17)), click-ready tags (DAz/DYn series of azido-/alkyne-analogs of DCP (19, 48–50); linear alkyne beta-ketoesters (18)), and more recently mitochondrial-targeting components (DCP-NEt_2_C and DCP-Rho1) (14, 16).

The goal of the work presented here was to build on the established selectivity of DCP-based reagents towards sulfenylated proteins (14, 16–18, 47), and on the clinical evidence of increased protein sulfenylation in radiation sensitive HNSCC tumors (10) to develop a first-generation PET radiotracer for imaging protein sulfenylation in vivo.

Thus, [^18^F]DCP and its cold analog [^19^F]DCP were synthesized and evaluated using recombinant model proteins (**Figure 1**), in vitro cell culture (**Figure 2, A-C**) and in vivo (**Figure 2, D-H, 3** and **4**). As described in the Results, the [^18^F]DCP shows very good serum stability (**Figure 2A**) and ability to distinguish between radiation sensitive and resistant cells both in cell culture (**Figure 2, A-C**) and in xenograft mouse models of HNSCC (**Figure 2, D-H**). Measurement of vascular parameters using quantitative optical spectroscopy exclude lower perfusion or hypoxia as determinants of decreased [^18^F]DCP incorporation signal in rSCC-61 tumors (**Figure 3**). The time course biodistribution analysis of [^18^F]DCP in SCC-61 and rSCC-61 xenograft mice (**Figure 2, D-H**) and the single time point analysis in an extended panel of radiation sensitive and resistant HNSCC tumors (**Figure 4**) show consistent tumor-to-muscle ratio and consistent 2-4 fold difference in radiotracer accumulation between the radiation sensitive and resistant tumors. The observed accumulation of [^18^F]DCP in organs such as the kidney and liver is expected given the metabolism and function of these organs. Selective tumor targeting can be achieved as a future goal with other means including the use of aptamers (51), liposomes (52), and others (53). EGF-liposomes could be particularly useful considering the 80-90% frequency of EGFR overexpression in HNSCC (54).

In conclusion, we have developed a novel PET radiotracer [^18^F]DCP for in vivo detection of protein sulfenylation capable of sensing differences in this oxidative modification between the radiation sensitive and resistant HNSCC cells in vitro and in animal models. We anticipate that such an ability to profile protein oxidation will contribute to accurate clinical assessment of tumor redox state aiding prognosis of response to radiation therapy before radiation treatment is attempted. This imaging agent could also be applied to identify patients in need of adjuvant redox altering therapy to improve clinical outcomes with radiation treatment.

## Materials and Methods

### Kinetic analysis of [^19^F]DCP reaction with C165A AhpC-SOH

Recombinant C165A mutant of *Salmonella typhimurium* AhpC was purified and oxidized as described in previous work (2, 14–18). The C165A AhpC-SOH (50 μM, also containing -SN and -SO_2/3_H species) was incubated with 5 mM dimedone or 5 mM [^19^F]DCP at room temperature in ammonium bicarbonate buffer pH 8.0 and analyzed at different time points (15 min, 30 min, 60 min, 120 min and 180 min) by ESI-TOF MS (Electrospray Ionization Time-of-Flight Mass Spectrometry) on an Agilent 6120 MSD-TOF system. The analysis was performed in positive ion mode with the following settings: capillary voltage: 3.5 kV, nebulizer gas pressure: 30 psig, drying gas flow: 5 L/min, fragmentor voltage: 200 V, skimmer voltage: 65 V, and gas temperature: 325 °C. Processing of data to generate the deconvoluted MS spectra was performed with Agilent MassHunter Workstation software v B.02.00. Kinetics analysis was performed by fitting the relative abundance of dimedone and [^19^F]DCP AhpC covalent adducts to an exponential equation in GraphPad Prism 7.0.

### Serum stability of [^18^F]DCP

The *ex vivo* serum stability of the radiotracer [^18^F]DCP was performed following previously published methods (55–57) and human serum (Sigma Aldrich). [^18^F]DCP (~1-2 mCi) was added to the serum to a final volume of 1 mL, and incubated at 37 °C. [^18^F]DCP and the radioactive serum mixture (~50 μL) were injected into a Quality Control High-Performance Liquid Chromatography (QC-HPLC) system at 5 min, 30 min, 1 h, 1.5 h, 2 h, 2.5 h, 3 h and 4 h post-radiotracer synthesis and serum incubation, respectively. There were no additional radiochemical species detected (e.g., resulting from defluorination of [^18^F] radiotracers (58–61)).

### In vitro cellular uptake assay

Human HNSCC SCC-61 and rSCC-61 cells extensively characterized in previous publications (10, 11, 22, 23) were utilized for the majority of the studies here. The SCC-61 cells were obtained from Dr. Ralph Weichselbaum, University of Chicago), and the rSCC-61 cells were generated *in vitro* by repeated irradiation and selection of a radiation resistant clone (10). Both SCC-61 and rSCC-61 cells were cultured in DMEM/F12 medium (Gibco) supplemented with 10% FBS (Gibco) and 1% Pen/Strep (Gibco) at 37°C and 5% CO_2_. Following PBS washing, cells were spun down, resuspended in growth media, and counted using a hemocytometer.

The *in vitro* [^18^F]DCP cellular uptake assays were performed following published protocols (55, 62, 63). Cells (1 × 10^5^) were seeded into each well of a 6-well culture and incubated overnight. On the day of the assay, fresh solution of non-radioactive compound (10 μM) in the respective cell medium was used as the blocker solution. The blocker solution was added 15 min prior to addition of radiotracer. Cells were then incubated with [^18^F]DCP (1 μCi/well) for 5 min, 30 min, 60 min (n=3) at 37 °C. Cellular uptake was allowed to proceed for selected time periods and then terminated by rinsing the cells with 1 mL of the ice-cold PBS. Residual fluid was removed by pipette, and 200 μL of 0.1% aqueous sodium dodecylsulfate (SDS) lysis solution was added to each well. The plate was then agitated at room temperature and 1 mL of the lysate was taken from each well for gamma counting. The radioactivity was counted using the Wallac 1480 Wizard gamma counter (Perkin Elmer). Additional 20 μL aliquots were taken in triplicate from each well for protein concentration determination using the Pierce bicinchoninic acid protein assay kit. The radioactivity data in each sample from each well were expressed as counts per minute (cpm) and were decay corrected for elapsed time. The cpm values of each well were normalized to the amount of radioactivity added to each well and the protein concentration in the well and expressed as percent uptake relative to the control condition. The data were expressed as percentage injection dose (%ID) per mg of protein present in each well with p values ≤ 0.05 considered statistically significant.

### Animal tumor implantation

Female athymic nude mice (Jackson Laboratory; average weight 18-20 grams, 6-7 weeks old) were housed in a pathogen-free facility of the Animal Research Program at Wake Forest University School of Medicine under a 12:12 h light/dark cycle and fed ad libitum. All animal experiments were conducted under IACUC approved protocols in compliance with the guidelines for the care and use of research animals established by Wake Forest University Medical School Animal Studies Committee (IACUC# A17-069). 1 - 2 × 10^6^ SCC-61, rSCC-61, JHU022 and SQ20B cells transfected with luciferase were suspended in 50 μL PBS and then mixed with 50 μL of Matrigel (BD Biosciences). The whole mixture was then injected subcutaneously in mice either in the right flank or between the shoulder blades. The presence of viable tumors was confirmed using bioluminescence imaging by injecting 150 mg luciferin/kg body weight (10 μL of 15 mg/mL luciferin/g of body weight) intra-peritoneally 15 min before imaging with IVIS Lumina LT Series III.

### Optical measurements and data analysis

These *in vivo* experiments were performed according to a protocol approved by the Animal Research Program at Wake Forest University School of Medicine (IACUC# A16-164). 1 × 10^6^ SCC-61 or rSCC-61 cells were suspended in 50 μL PBS and then mixed with 50 μL of Matrigel (BD Biosciences). The whole mixture was then injected in the right flank of the mice subcutaneously. The size of viable tumors was estimated using a caliper and the formula V = (W^2^ x L)/2 where V is tumor volume, W is tumor width, and L is tumor length. Metabolic imaging was performed 1-2 weeks after tumor implantation. The mice were fasted for 6 hours prior to each set of optical experiments to maintain minimal variance in metabolic demand among animals (24). All measured mice received a tail-vein injection of TMRE (100 μL of 75 μM) first and then a tail-vein injection 2-NBDG (100 μL of 6 mM 2-NBDG) with 20 minute delay as reported previously (25, 64). The 2-NBDG uptake at 60 minutes post 2-NBDG injection and TMRE uptake at 80 minutes post TMRE injection were used to report final 2-NBDG and TMRE uptake.

A quantitative optical spectroscopy system introduced previously (24, 25) was used to perform diffuse reflectance and fluorescence measurements in a darkroom. Optical measurements on animals were performed by placing the fiber probe gently on the tumors and fixing the measurement on the center of the tumor mass (25). The measured spot size was 2.0-mm in diameter which was determined by the probe size as reported previously (25, 64). Baseline diffuse reflectance and fluorescence spectra were measured from the tumor sites prior to any probe injection. Diffuse reflectance spectra were acquired from 450 nm to 650 nm with an integration time of 3.8 ms. The 2-NDBG fluorescence spectra between 520 nm to 600 nm were measured with an excitation light of 488 nm and an integration time of 2 seconds. The TMRE fluorescence spectra between 565 nm to 650 nm were obtained with an excitation light of 555 nm and an integration time of 5 seconds. Optical measurements on each animal were acquired continuously for a period of 80 minutes. All diffuse reflectance spectra were calibrated using a reflectance puck, while all fluorescence spectra were calibrated with fluorescence standard puck (24, 25). All optical spectral data were processed using the previously developed inverse Monte Carlo (MC) model (24) to extract tissue absorption spectra, tissue background distortion free fluorescence of 2-NBDG, and TMRE (25). The extracted absorption spectra were processed to estimate oxygen saturation and total hemoglobin concentration [Hb] (65). The optical measurements at the last time point were used to calculate final 2-NBDG or TMRE uptake.

### MicroPET/CT imaging

MicroPET/CT imaging experiments were performed in SCC-61, JHU022, SQ20B, and rSCC-61 tumor bearing mice (n = 4, 25 ± 2.5 g) using Trifoil PET/CT scanner (1.2 mm resolution and 14 cm axial field of view). Mice were placed in an induction chamber containing ~2% isoflurane/oxygen then secured to a custom double bed for placement of tail vein catheters; anesthesia was maintained via nose-cone at ~1% isoflurane/oxygen throughout the scanning. [^18^F]DCP was intravenously injected and scanned 50 min post-injection. A 3 min CT scan was obtained prior to the PET scan. Four SCC-61 tumor bearing mice from the same cohort were used for blocking experiments. These mice received 10 mg/kg 100 μL of the non-radioactive [^19^F]DCP, 30 min prior to the injection of radiotracer. The images were reconstructed using TriFoil attenuation correction and Fourier rebinning parameters (56, 66). The reconstructed CT and PET images were co-registered and analyzed using the TriFoil filtered back-projection 3D-OSEM algorithm.

### Biodistribution analysis

Biodistribution experiments were performed first in SCC-61 and rSCC-61 tumor bearing mice (n = 4, 25 ± 2.5 g). Mice were placed in an induction chamber containing ~2% isoflurane/oxygen and then secured to a custom bed for placement of tail vein catheters. [^18^F]DCP (60-80 μCi in ~65-80 μL saline containing 10% ethanol, with a specific activity of 2610-2820 mCi/μmol) was administered through tail vein injection. Mice were sacrificed by decapitation via cervical dislocation under anesthesia at 5, 30, 60, 90 and 120 min (n = 4 for each time point). For all other tumor mice biodistribution studies (e.g., Figure 4: SCC-61, JHU022, SQ20B, and rSCC-61), mice were sacrificed at 60 min post radiotracer injection. The specimens of interest including blood, brain, tumor, heart, lungs, liver, spleen, kidneys, pancreas, muscle and bone were removed, weighed, and counted in an automatic gamma counter (Wallac 2480 Wizard). Radiotracer uptake was calculated as percentages of injected dose per gram of tissue (%ID/g tissue).

### Statistical analysis

Statistical analysis (t-test) was performed using Microsoft Excel and all results are displayed as mean ± standard deviation. Asterisks indicate statistically significant changes [α = 0.05, P values of 0.01–0.05 (*), 0.001–0.01 (**), or <0.001 (***)].

## Author contributions

C.M.F. developed the overall concept for the work presented here. Z.L. and S.A.V. performed chemical synthesis of [^19^F]DCP and cold precursors of [^18^F]DCP under the supervision of S.B.K. X.C., K.S., T.E.F. and H.W. performed all other experiments under the supervision of C.M.F., A.W.T., and K.K.S.S. Optical measurements and data processing were performed by C.Z. with support from M.M. under the supervision of N.M. and M.D. X.C. provided the animals for these studies. L.B.P. provided critical intellectual feedback for the overall project.

## Competing interests

C.M.F., S.B.K., L.B.P., and K.K.S.S. filed a provisional patent application for the use of protein sulfenylation probes like the [^18^F]DCP described here for PET imaging and chemotherapeutic applications. C.M.F. S.B.K. and L.B.P. are co-founders of Xoder Technologies, LLC.

## Data and materials availability

Not applicable.

## Acknowledgments

The authors thank Kimbrell family for the support of high-end mass spectrometry instrumentation in C.M.F.’ s laboratory.

## Funding

The authors acknowledge financial support for these studies provided by the Center for Redox Biology and Medicine at Wake Forest School of Medicine (pilot funds to S.B.K.), the Wake Forest Baptist Comprehensive Cancer Center (NIH/NCI P30 CA12197; pilot funds to C.M.F. and support of shared resources facilities), and NIH/NCI R42CA156901 to N.R..

## Notes

### Competing Interest Statement

The authors have declared no competing interest.

## References

1. Garcia-Peris P, Paron L, Velasco C, de la Cuerda C, Camblor M, Breton I, et al. Long-term prevalence of oropharyngeal dysphagia in head and neck cancer patients: Impact on quality of life. Clin Nutr. 2007;26(6):710–7.

2. Reisz JA, Bansal N, Qian J, Zhao W, and Furdui CM. Effects of ionizing radiation on biological molecules--mechanisms of damage and emerging methods of detection. Antioxid Redox Signal. 2014;21(2):260–92.

3. Anderson CM, Lee CM, Saunders DP, Curtis A, Dunlap N, Nangia C, et al. Phase IIb, Randomized, Double-Blind Trial of GC4419 Versus Placebo to Reduce Severe Oral Mucositis Due to Concurrent Radiotherapy and Cisplatin For Head and Neck Cancer. J Clin Oncol. 2019;37(34):3256–65.

4. Anderson CM, Sonis ST, Lee CM, Adkins D, Allen BG, Sun W, et al. Phase 1b/2a Trial of the Superoxide Dismutase Mimetic GC4419 to Reduce Chemoradiotherapy-Induced Oral Mucositis in Patients With Oral Cavity or Oropharyngeal Carcinoma. Int J Radiat Oncol Biol Phys. 2018;100(2):427–35.

5. Birer SR, Lee CT, Choudhury KR, Young KH, Spasojevic I, Batinic-Haberle I, et al. Inhibition of the Continuum of Radiation-Induced Normal Tissue Injury by a Redox-Active Mn Porphyrin. Radiat Res. 2017;188(1):94–104.

6. Brizel DM, Wasserman TH, Henke M, Strnad V, Rudat V, Monnier A, et al. Phase III randomized trial of amifostine as a radioprotector in head and neck cancer. J Clin Oncol. 2000;18(19):3339–45.

7. Garcia MK, Meng Z, Rosenthal DI, Shen Y, Chambers M, Yang P, et al. Effect of True and Sham Acupuncture on Radiation-Induced Xerostomia Among Patients With Head and Neck Cancer: A Randomized Clinical Trial. JAMA Netw Open. 2019;2(12):e1916910.

8. Purandare NC, Puranik AD, Shah S, Agrawal A, and Rangarajan V. Post-treatment appearances, pitfalls, and patterns of failure in head and neck cancer on FDG PET/CT imaging. Indian J Nucl Med. 2014;29(3):151–7.

9. Castaldi P, Leccisotti L, Bussu F, Micciche F, and Rufini V. Role of (18)F-FDG PET-CT in head and neck squamous cell carcinoma. Acta Otorhinolaryngol Ital. 2013;33(1):1–8.

10. Bansal N, Mims J, Kuremsky JG, Olex AL, Zhao W, Yin L, et al. Broad phenotypic changes associated with gain of radiation resistance in head and neck squamous cell cancer. Antioxid Redox Signal. 2014;21(2):221–36.

11. Mims J, Bansal N, Bharadwaj MS, Chen X, Molina AJ, Tsang AW, et al. Energy metabolism in a matched model of radiation resistance for head and neck squamous cell cancer. Radiat Res. 2015;183(3):291–304.

12. Devarie-Baez NO, Silva Lopez EI, and Furdui CM. Biological chemistry and functionality of protein sulfenic acids and related thiol modifications. Free Radic Res. 2016;50(2):172–94.

13. Poole LB. The basics of thiols and cysteines in redox biology and chemistry. Free Radic Biol Med. 2015;80:148–57.

14. Holmila RJ, Vance SA, Chen X, Wu H, Shukla K, Bharadwaj MS, et al. Mitochondria-targeted Probes for Imaging Protein Sulfenylation. Sci Rep. 2018;8(1):6635.

15. Klomsiri C, Nelson KJ, Bechtold E, Soito L, Johnson LC, Lowther WT, et al. Use of dimedone-based chemical probes for sulfenic acid detection evaluation of conditions affecting probe incorporation into redox-sensitive proteins. Methods Enzymol. 2010;473:77–94.

16. Poole LB, Klomsiri C, Knaggs SA, Furdui CM, Nelson KJ, Thomas MJ, et al. Fluorescent and affinity-based tools to detect cysteine sulfenic acid formation in proteins. Bioconjug Chem. 2007;18(6):2004–17.

17. Qian J, Klomsiri C, Wright MW, King SB, Tsang AW, Poole LB, et al. Simple synthesis of 1,3-cyclopentanedione derived probes for labeling sulfenic acid proteins. Chem Commun (Camb). 2011;47(32):9203–5.

18. Qian J, Wani R, Klomsiri C, Poole LB, Tsang AW, and Furdui CM. A simple and effective strategy for labeling cysteine sulfenic acid in proteins by utilization of beta-ketoesters as cleavable probes. Chem Commun (Camb). 2012;48(34):4091–3.

19. Bechtold E. Department of Chemistry. Wake Forest University; 2010.

20. Demko ZP, and Sharpless KB. An intramolecular [2 + 3] cycloaddition route to fused 5-heterosubstituted tetrazoles. Org Lett. 2001;3(25):4091–4.

21. Bai S, Li S, Xu J, Peng X, Sai K, Chu W, et al. Synthesis and structure-activity relationship studies of conformationally flexible tetrahydroisoquinolinyl triazole carboxamide and triazole substituted benzamide analogues as sigma2 receptor ligands. J Med Chem. 2014;57(10):4239–51.

22. Chen X, Liu L, Mims J, Punska EC, Williams KE, Zhao W, et al. Analysis of DNA methylation and gene expression in radiation-resistant head and neck tumors. Epigenetics. 2015;10(6):545–61.

23. Lewis JE, Costantini F, Mims J, Chen X, Furdui CM, Boothman DA, et al. Genome-Scale Modeling of NADPH-Driven beta-Lapachone Sensitization in Head and Neck Squamous Cell Carcinoma. Antioxid Redox Signal. 2018;29(10):937–52.

24. Rajaram N, Reesor AF, Mulvey CS, Frees AE, and Ramanujam N. Non-invasive, simultaneous quantification of vascular oxygenation and glucose uptake in tissue. PLoS One. 2015;10(1):e0117132.

25. Zhu C, Martin HL, Crouch BT, Martinez AF, Li M, Palmer GM, et al. Near-simultaneous quantification of glucose uptake, mitochondrial membrane potential, and vascular parameters in murine flank tumors using quantitative diffuse reflectance and fluorescence spectroscopy. Biomed Opt Express. 2018;9(7):3399–412.

26. Flach S, Maniam P, and Manickavasagam J. E-cigarettes and head and neck cancers: A systematic review of the current literature. Clin Otolaryngol. 2019;44(5):749–56.

27. Nocini R, Lippi G, and Mattiuzzi C. The worldwide burden of smoking-related oral cancer deaths. Clin Exp Dent Res. 2020;6(2):161–4.

28. Ang KK, Harris J, Wheeler R, Weber R, Rosenthal DI, Nguyen-Tan PF, et al. Human papillomavirus and survival of patients with oropharyngeal cancer. N Engl J Med. 2010;363(1):24–35.

29. Fakhry C, Westra WH, Li S, Cmelak A, Ridge JA, Pinto H, et al. Improved survival of patients with human papillomavirus-positive head and neck squamous cell carcinoma in a prospective clinical trial. J Natl Cancer Inst. 2008;100(4):261–9.

30. Brieger K, Schiavone S, Miller FJ, Jr., and Krause KH. Reactive oxygen species: from health to disease. Swiss Med Wkly. 2012;142:w13659.

31. Sabharwal SS, and Schumacker PT. Mitochondrial ROS in cancer: initiators, amplifiers or an Achilles’ heel? Nat Rev Cancer. 2014;14(11):709–21.

32. Liou GY, and Storz P. Reactive oxygen species in cancer. Free Radic Res. 2010;44(5):479–96.

33. Pelicano H, Carney D, and Huang P. ROS stress in cancer cells and therapeutic implications. Drug Resist Updat. 2004;7(2):97–110.

34. Reczek CR, and Chandel NS. The Two Faces of Reactive Oxygen Species in Cancer. Annual Review of Cancer Biology. 2016.

35. Gorrini C, Harris IS, and Mak TW. Modulation of oxidative stress as an anticancer strategy. Nat Rev Drug Discov. 2013;12(12):931–47.

36. Damia G, and Garattini S. The pharmacological point of view of resistance to therapy in tumors. Cancer Treat Rev. 2014;40(8):909–16.

37. Holohan C, Van Schaeybroeck S, Longley DB, and Johnston PG. Cancer drug resistance: an evolving paradigm. Nat Rev Cancer. 2013;13(10):714–26.

38. Maiti AK. Reactive Oxygen Species Reduction is a Key Underlying Mechanism of Drug Resistance in Cancer Chemotherapy. Chemotherapy. 2012.

39. Singer E, Judkins J, Salomonis N, Matlaf L, Soteropoulos P, McAllister S, et al. Reactive oxygen species-mediated therapeutic response and resistance in glioblastoma. Cell Death Dis. 2015;6:e1601.

40. Diehn M, Cho RW, Lobo NA, Kalisky T, Dorie MJ, Kulp AN, et al. Association of reactive oxygen species levels and radioresistance in cancer stem cells. Nature. 2009;458(7239):780–3.

41. Singh A, Bodas M, Wakabayashi N, Bunz F, and Biswal S. Gain of Nrf2 function in non-small-cell lung cancer cells confers radioresistance. Antioxid Redox Signal. 2010;13(11):1627–37.

42. Hwang IT, Chung YM, Kim JJ, Chung JS, Kim BS, Kim HJ, et al. Drug resistance to 5-FU linked to reactive oxygen species modulator 1. Biochem Biophys Res Commun. 2007;359(2):304–10.

43. Carroll VN, Truillet C, Shen B, Flavell RR, Shao X, Evans MJ, et al. [(11)C]Ascorbic and [(11)C]dehydroascorbic acid, an endogenous redox pair for sensing reactive oxygen species using positron emission tomography. Chem Commun (Camb). 2016;52(27):4888–90.

44. Qin H, Carroll VN, Sriram R, Villanueva-Meyer JE, von Morze C, Wang ZJ, et al. Imaging glutathione depletion in the rat brain using ascorbate-derived hyperpolarized MR and PET probes. Sci Rep. 2018;8(1):7928.

45. Solingapuram Sai KK, Bashetti N, Chen X, Norman S, Hines JW, Meka O, et al. Initial biological evaluations of (18)F-KS1, a novel ascorbate derivative to image oxidative stress in cancer. EJNMMI Res. 2019;9(1):43.

46. Tsujikawa T, Asahi S, Oh M, Sato Y, Narita N, Makino A, et al. Assessment of the Tumor Redox Status in Head and Neck Cancer by 62Cu-ATSM PET. PLoS One. 2016;11(5):e0155635.

47. Poole LB, Zeng BB, Knaggs SA, Yakubu M, and King SB. Synthesis of chemical probes to map sulfenic acid modifications on proteins. Bioconjug Chem. 2005;16(6):1624–8.

48. Reddie KG, Seo YH, Muse Iii WB, Leonard SE, and Carroll KS. A chemical approach for detecting sulfenic acid-modified proteins in living cells. Mol Biosyst. 2008;4(6):521–31.

49. Seo YH, and Carroll KS. Facile synthesis and biological evaluation of a cell-permeable probe to detect redox-regulated proteins. Bioorganic & Medicinal Chemistry Letters. 2009;19(2):356–9.

50. Truong TH, Garcia FJ, Seo YH, and Carroll KS. Isotope-coded chemical reporter and acid-cleavable affinity reagents for monitoring protein sulfenic acids. Bioorganic & Medicinal Chemistry Letters. 2011;21(17):5015–20.

51. Barbas AS, Mi J, Clary BM, and White RR. Aptamer applications for targeted cancer therapy. Future Oncol. 2010;6(7):1117–26.

52. Zalba S, Contreras AM, Merino M, Navarro I, de Ilarduya CT, Troconiz IF, et al. EGF-liposomes promote efficient EGFR targeting in xenograft colocarcinoma model. Nanomedicine (Lond). 2016;11(5):465–77.

53. Raut S, Mooberry L, Sabnis N, Garud A, Dossou AS, and Lacko A. Reconstituted HDL: Drug Delivery Platform for Overcoming Biological Barriers to Cancer Therapy. Front Pharmacol. 2018;9:1154.

54. Harari PM, Wheeler DL, and Grandis JR. Molecular target approaches in head and neck cancer: epidermal growth factor receptor and beyond. Semin Radiat Oncol. 2009;19(1):63–8.

55. Sai KKS, Sattiraju A, Almaguel FG, Xuan A, Rideout S, Krishnaswamy RS, et al. Peptide-based PET imaging of the tumor restricted IL13RA2 biomarker. Oncotarget. 2017;5.

56. Sattiraju A, Pandya D, Solingapuram Sai K, Wadas T, Herpai D, Debinski W, et al. IL13RA2 targeted alpha particle therapy against glioblastoma. Journal of Nuclear Medicine. 2016;57(supplement 2):634.

57. Wang H-E, Wu S-Y, Chang C-W, Liu R-S, Hwang L-C, Lee T-W, et al. Evaluation of F-18-labeled amino acid derivatives and [18F]FDG as PET probes in a brain tumor-bearing animal model. Nuclear Medicine and Biology. 2005;32(4):367–75.

58. Harada R, Furumoto S, Tago T, Katsutoshi F, Ishiki A, Tomita N, et al. Characterization of the radiolabeled metabolite of tau PET tracer 18F-THK5351. European Journal of Nuclear Medicine and Molecular Imaging. 2016;43(12):2211–8.

59. Jin H, Yue X, Zhang X, Li J, Yang H, Flores H, et al. A promising F-18 labeled PET radiotracer (-)-[18F]VAT for assessing the VAChT in vivo. Journal of Nuclear Medicine. 2015;56(supplement 3):4.

60. Koglin N, Friebe M, Berndt M, Graham K, Krasikova R, Kuznetsova O, et al. [F-18]BAY 85-8050: A novel tumor specific probe for PET imaging - Preclinical results. Journal of Nuclear Medicine. 2010;51(supplement 2):1535.

61. Pacelli A, Greenman J, Cawthorne C, and Smith G. Imaging COX-2 expression in cancer using PET/SPECT radioligands: current status and future directions. Journal of Labelled Compounds and Radiopharmaceuticals. 2014;57(4):317–22.

62. Yamamoto Y, Nishiyama Y, Ishikawa S, Nakano J, Chang SS, Bandoh S, et al. Correlation of 18F-FLT and 18F-FDG uptake on PET with Ki-67 immunohistochemistry in non-small cell lung cancer. European Journal of Nuclear Medicine and Molecular Imaging. 2007;34(10):1610–6.

63. Solingapuram Sai KK, Das BC, Sattiraju A, Almaguel FG, Craft S, and Mintz A. Radiolabeling and initial biological evaluation of [18F]KBM-1 for imaging RAR-α receptors in neuroblastoma. Bioorganic & Medicinal Chemistry Letters. 2017;27(6):1425–7.

64. Zhu C, Martinez AF, Martin HL, Li M, Crouch BT, Carlson DA, et al. Near-simultaneous intravital microscopy of glucose uptake and mitochondrial membrane potential, key endpoints that reflect major metabolic axes in cancer. Sci Rep. 2017;7(1):13772.

65. Franceschini MA, Gratton E, and Fantini S. Noninvasive optical method of measuring tissue and arterial saturation: an application to absolute pulse oximetry of the brain. Opt Lett. 1999;24(12):829–31.

66. Solingapuram Sai KK, Prabhakaran J, Ramanathan G, Rideout S, Whitlow C, Mintz A, et al. Radiosynthesis and Evaluation of [(11)C]HD-800, a High Affinity Brain Penetrant PET Tracer for Imaging Microtubules. ACS Med Chem Lett. 2018;9(5):452–6.

