## Supplemental Material for "[^18^F]DCP, First Generation PET Radiotracer for Diagnosis of Radiation Resistant Head and Neck Cancer"

**Supplementary Materials and Methods**

1. Chemical Synthesis of [$^{18}$F]DCP Cold Precursors

2. Radiochemical Synthesis of [$^{18}$F]DCP from Precursor 2

3. Chemical Synthesis of cold [$^{19}$F]DCP

**Supplementary Figures**

Figure S1. Toluenesulfonic acid-2-azidoethyl ester. (**A**) $^{1}$H-NMR spectrum. (**B**) $^{13}$C-NMR spectrum.

Figure S2. Characterization of precursor **1** in [$^{18}$F]DCP synthesis. (**A**) $^{1}$H-NMR spectrum. (**B**) $^{13}$C-NMR spectrum.

Figure S3. Characterization of precursor **2** in [$^{18}$F]DCP synthesis. (**A**) $^{1}$H-NMR spectrum. (**B**) $^{13}$C-NMR spectrum.

Figure S4. Quality control analysis of [$^{18}$F]DCP showing consistent chromatographic properties with [$^{19}$F]DCP.

Figure S5. Characterization of precursor **3** in cold [$^{19}$F]DCP synthesis. (**A**) $^{1}$H-NMR spectrum. (**B**) $^{13}$C-NMR spectrum.

Figure S6. Characterization of cold [$^{19}$F]DCP. (**A**) $^{1}$H-NMR spectrum. (**B**) $^{13}$C-NMR spectrum. (**C**) $^{19}$F-NMR spectrum.

### 1. Chemical Synthesis of [$^{18}$F]DCP Cold Precursors

**General**. Reagents were obtained from commercial sources and used without additional purification. Reaction solvents such as dichloromethane, acetonitrile, and ethyl acetate were dried and distilled over calcium hydride. Extraction, silica, and preparative reverse phase chromatography solvents were technical grade. Analytical TLC was performed on silica gel plates, and visualization was accomplished with UV light. $^{1}$H NMR spectra were recorded on Bruker Avance DPX-300 and DRX-500 instruments at 300.13 and 500.13 MHz, respectively. $^{13}$C NMR spectra were recorded on the described instruments operating at 75.48 and 125.76 MHz, respectively. $^{19}$F NMR spectra were recorded on the described instruments operating at 121.49 MHz and 202.46 MHz, respectively. Low-resolution mass spectra were obtained using an Agilent Technologies 1100 LC/MSD ion trap mass spectrometer equipped with an atmospheric pressure electrospray ionization source and was operated in positive ion mode unless otherwise noted.

**Toluenesulfonic acid-2-azidoethyl ester**

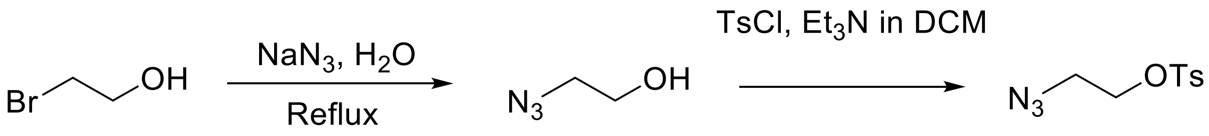


A mixture of 2-bromoethanol (6.761 g, 50 mmol) and sodium azide (3.915 g, 60 mmol) in water (15 mL) was heated at reflux in a 250 mL round bottom flask equipped with a magnetic stir bar. After 16 hours, the mixture was cooled to room temperature, saturated with $MgSO_4$ and extracted with dichloromethane (2 x 20 mL). The organic layers were combined, dried over anhydrous $MgSO_4$, filtered and the filtrate transferred to a clean 250 mL round bottom flask. Trimethylamine (10 mL) and toluene sulfonyl chloride (9.828 g, 50 mmol) were added to the filtrate and the resulting mixture was stirred at room temperature for 4 hours at which time glycine (0.758 g, 10 mmol) was added. After 2 more hours stirring at room temperature, the organic layer was washed with aqueous NaOH (1M, 2 x 100 mL), dried with anhydrous $MgSO_4$ and the solvent was removed under vacuum to yield a yellow oil (10.133 g) that was purified by silica gel column chromatography (eluent: petroleum ether: EtOAc = 3:1) to yield a light yellow oil (8.081 g, 33 mmol, 66 % yield).

**Data:**

**$^{1}$H-NMR ($CDCl_3$) δ:** 7.82 (d, $J$ = 9.0 Hz, 2H), 7.36 (d, $J$ = 9.0 Hz, 2H), 4.16 (t, $J$ = 4.5 Hz, 2H), 3.48 (t, $J$ = 4.5 Hz, 2H), 2.46 (s, 3H) (**Figure S1A**).

**$^{13}$C-NMR ($CDCl_3$) δ:** 145.28, 132.58, 130.00, 127.96, 68.11, 49.59, 21.68 (**Figure S1B**).

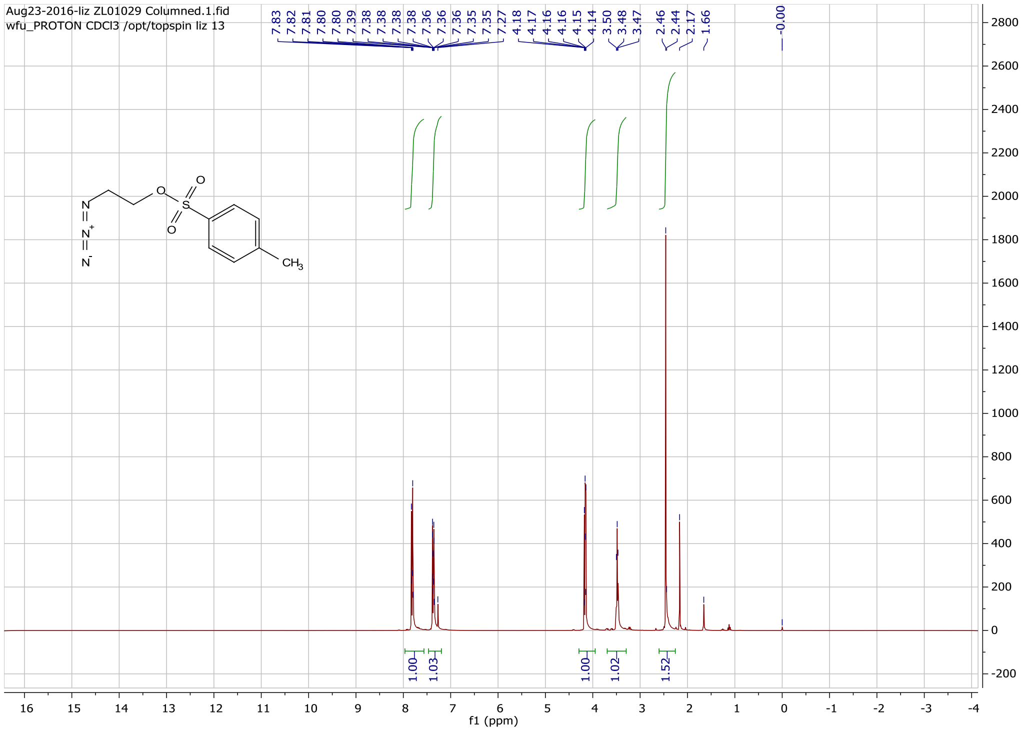


**Figure S1A. Toluenesulfonic acid-2-azidoethyl ester, $^1$H-NMR spectrum.**

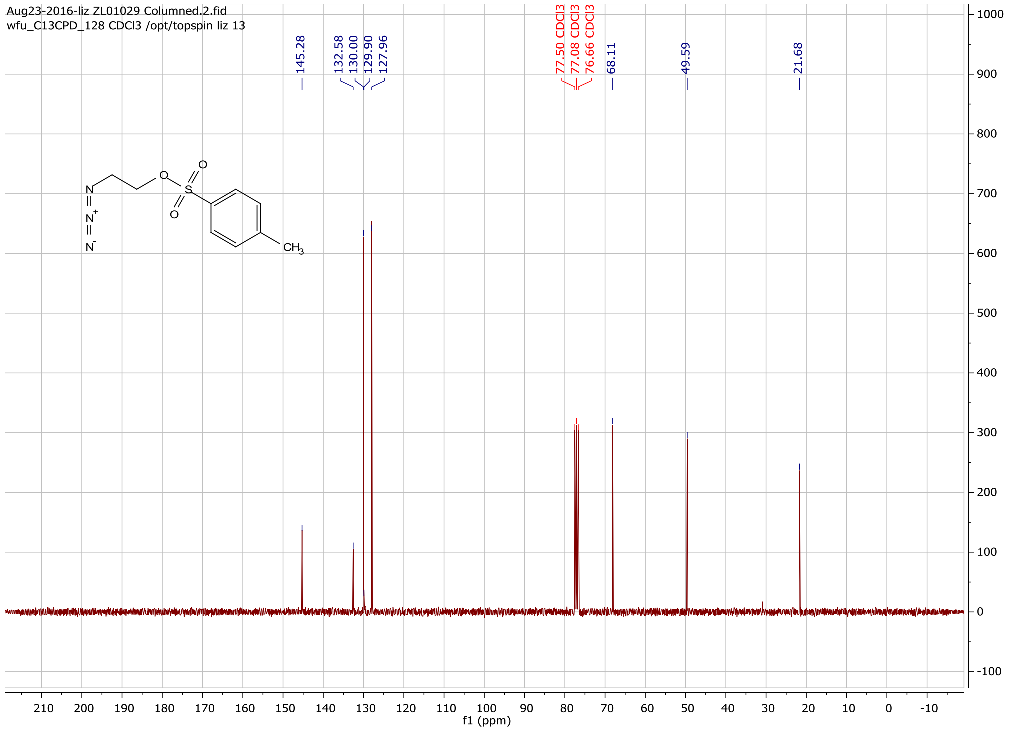


**Figure S1B. Toluenesulfonic acid-2-azidoethyl ester, $^{13}$C-NMR spectrum.**

**2-(4-(3-(4-ethoxy-2-oxocyclohex-3-en-1-yl)propyl)-*1H*-1,2,3-triazol-1-yl)ethyl 4-methylbenzenesulfonate (1)**

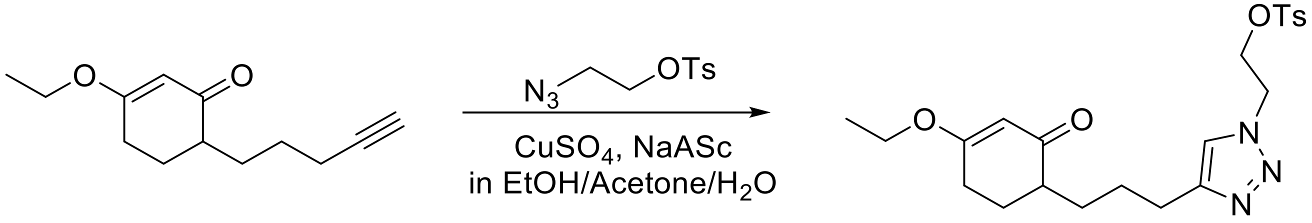


Sodium ascorbate (24 mg, 0.12 mmol) and $CuSO_4$ (16 µl of 1M solution in water) were added in sequence to a solution of 3-ethoxy-6-(pent-4-yn-1-yl)cyclohex-2-en-1-one (0.105 g, 0.5 mmol) and toluenesulfonic acid-2-azidoethyl ester (0.244 g, 1.0 mmol) in acetone (1 mL), ethanol (2 mL), and water (1 mL). After stirring for 3 days, TLC indicated the formation of a new product and the solvent was removed under vacuum to give a crude product that was purified by silica gel column chromatography (eluent: EtOAc) to give a sticky yellow solid (0.168 g, 0.37 mmol. 74 % yield).

**Data:**

**$^{1}$H-NMR ($CDCl_3$) δ:** 7.60 (d, *J* = 9.0 Hz, 2H), 7.31 (s, 1H), 7.25 (d, *J* = 9.0 Hz, 2H), 5.22 (s, 1H), 4.50 (t, *J* = 4.5 Hz, 2H), 4.31 (t, *J* = 4.5 Hz, 2H), 3.80 (q, *J* = 6.0 Hz, 2H), 2.67 – 2.56 (m, 2H), 2.36 – 2.29 (m, 6H), 2.18 – 2.09 (m, 1H), 2.04 – 1.95 (m, 1H), 1.85 – 1.76 (m, 1H), 1.72 – 1.58 (m, 3H), 1.44 – 1.18 (m, 2H), 1.27 (t, *J* = 6.0 Hz, 3H) (**Figure S2A**).

**$^{13}$C-NMR ($CDCl_3$) δ:** 201.34, 176.90, 148.07, 145.40, 132.04, 130.03, 127.77, 121.86, 102.10, 67.86, 64.18, 48.79, 44.89, 29.09, 28.05, 26.75, 26.26, 25.62, 21.64, 14.12 (**Figure S2B**).

**LRMS-ESI$^{+}$ (m/z):** $[M + H]^{+}$ predicted 448.2, found 448.2.

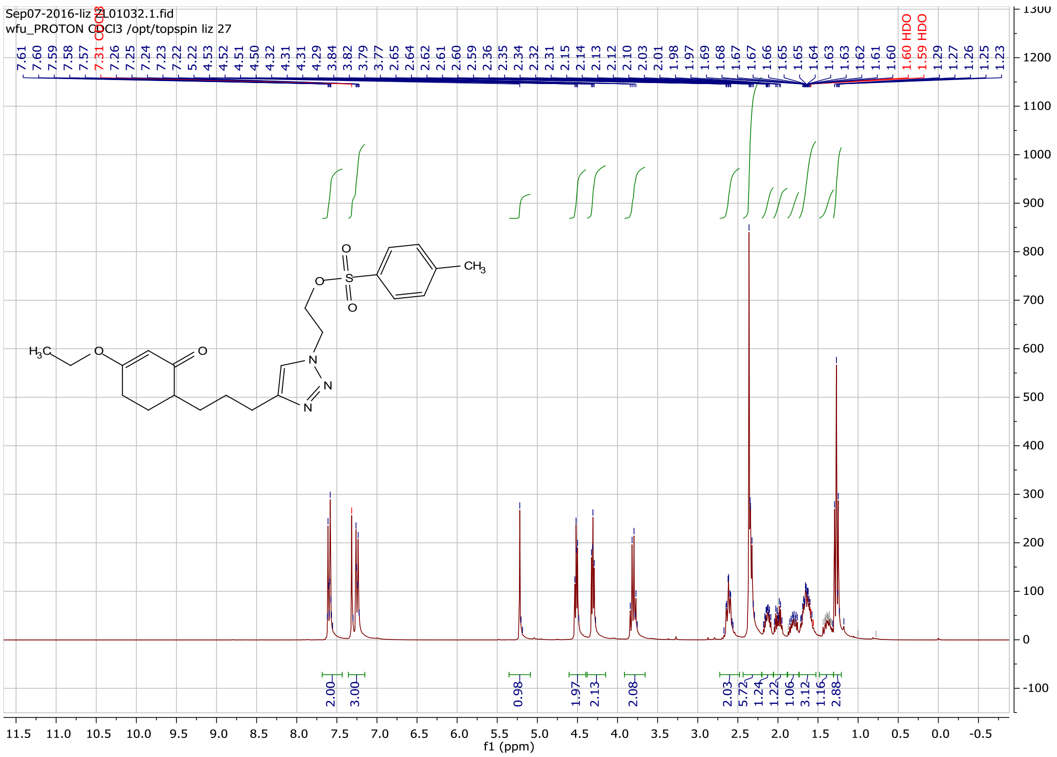

**Figure S2A. $^{1}$H-NMR spectrum of precursor 1 in [$^{18}$F]DCP synthesis.**

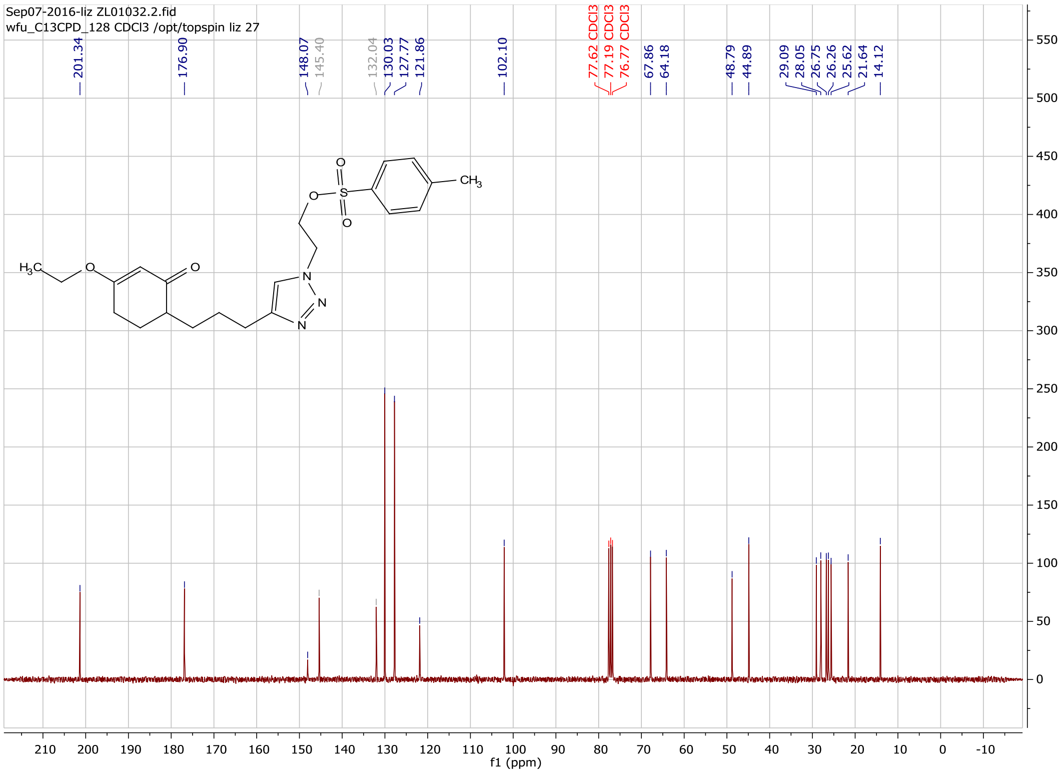

**Figure S2B. $^{13}$C-NMR spectrum of precursor 1 in [$^{18}$F]DCP synthesis.**

**2-(4-(3-(2,4-dioxocyclohexyl)propyl)-*1H*-1,2,3-triazol-1-yl)ethyl 4-methylbenzenesulfonate (2).**

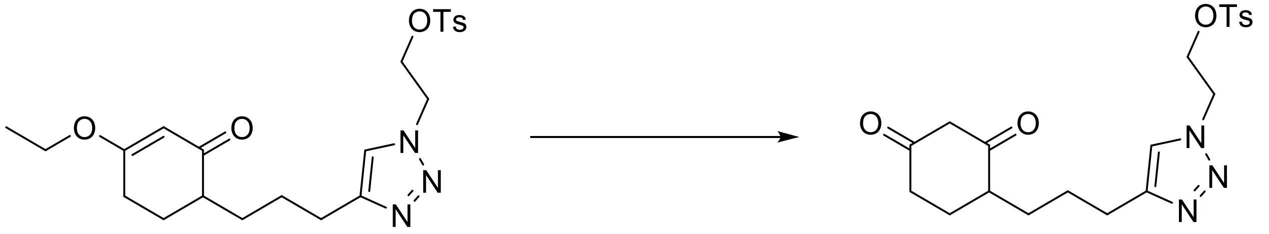


Ceric ammonium nitrate (CAN, 25 mg, 0.046 mmol) was added to a solution of 2-(4-(3-(4-ethoxy-2-oxocyclohex-3-en-1-yl)propyl)-*1H*-1,2,3-triazol-1-yl)ethyl 4-methylbenzenesulfonate (0.168 g, 0.37 mmol) in acetonitrile and water (3 mL : 3 mL) and the resulting solution was refluxed for 5 hours. Upon cooling, the solvent was removed under vacuum and the crude product purified by C18 reversed phase silica gel column chromatography (elution: acetonitrile in water, 3% to 50%). Lyophilization of the appropriate fractions provided a light-yellow powder (0.113 g, 0.27 mmol, 73 % yield).

**Data:**

**$^1$H-NMR ($CDCl_3$) δ:** 7.62 (d, $J$ = 9.0 Hz, 2H), 7.33 (s, 1H), 7.26 (d, $J$ = 9.0 Hz, 2H), 4.53 (t, $J$ = 4.5 Hz, 2H), 4.29 (t, $J$ = 4.5 Hz, 2H), 3.35 – 3.34 (m, 1H), 2.68 – 2.64 (m, 2H), 2.61 – 2.41 (m, 2H), 2.37 (s, 3H), 2.12 – 2.04 (m, 2H), 1.89 – 1.79 (m, 1H), 1.75 – 1.67 (m, 3H), 1.57 – 1.34 (m, 2H) (**Figure S3A**).

**$^{13}$C-NMR ($CDCl_3$) δ:** 204.60, 203.92, 147.76, 145.54, 132.09, 130.10, 127.85, 122.06, 67.84, 58.31, 49.10, 49.03, 39.65, 28.59, 26.66, 25.47, 24.48, 21.70 (**Figure S3B**).

**LRMS-ESI$^+$ (m/z):** $[M + H]^+$ predicted 420.2, found 420.1.

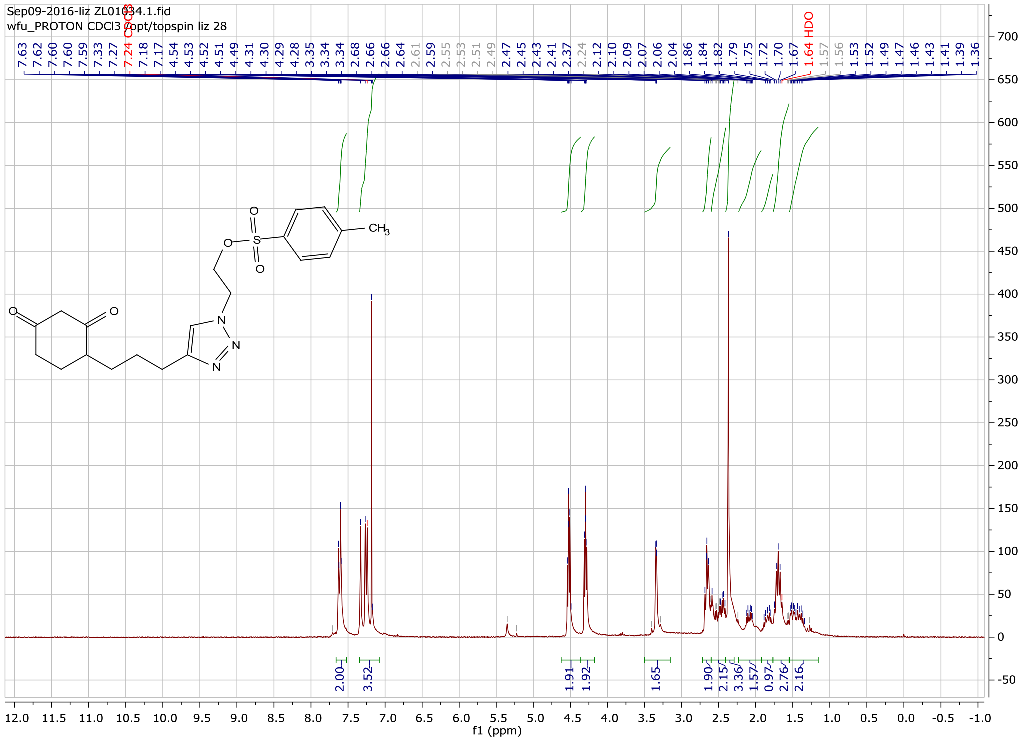

**Figure S3A.** $^{1}$**H-NMR spectrum of precursor 2 in [**$^{18}$**F]DCP synthesis.**

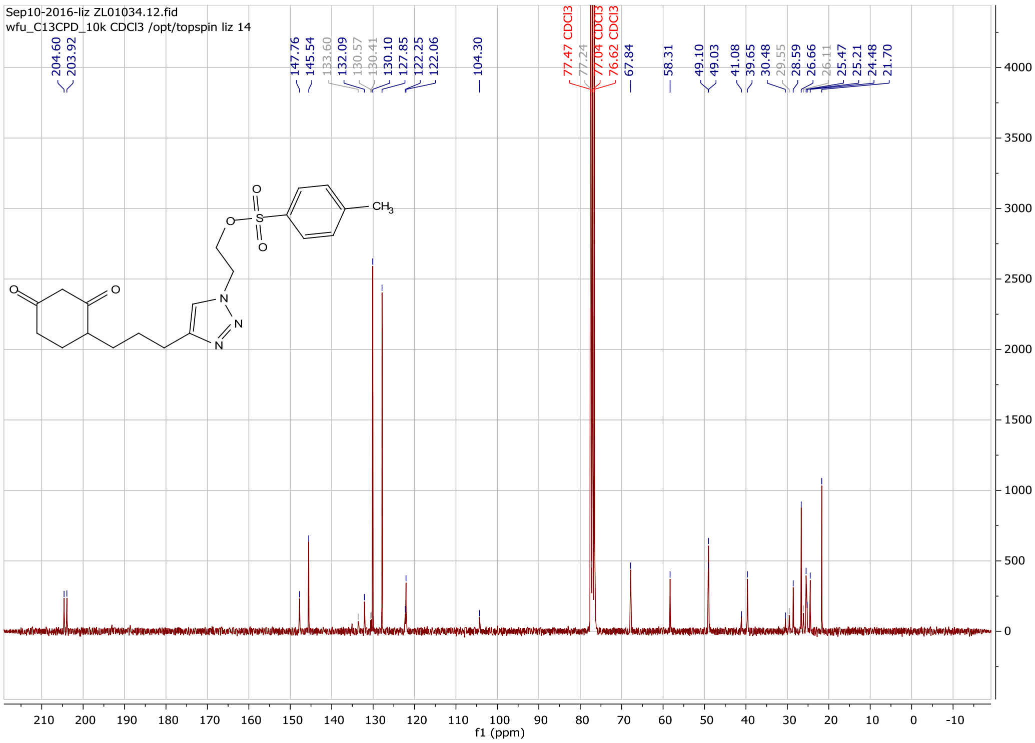

**Figure S3B.** $^{13}$**C-NMR spectrum of precursor 2 in [**$^{18}$**F]DCP synthesis.**

### 2. Radiochemical Synthesis of [$^{18}$F]DCP from Precursor 2

**Materials.** Anhydrous acetonitrile (ACN), anhydrous dimethylformamide (DMF), Kryptofix 222, HPLC grade $H_2O$, ammonium formate were purchased from Sigma-Aldrich and were used without any additional purification. All reactions were carried out using anhydrous solvents unless otherwise stated. C18 Sep Pak cartridge was purchased from Waters. Analytical HPLC was performed using Varian ProStar system, which includes quaternary gradient pump, manual injector, a variable wavelength detector and a standard Bioscan radioactivity-HPLC-flow detector. Both the HPLC columns i.e., semiprep HPLC- Phenomenex Luna C18 HPLC column: 250 x 10 mm, 10 µm and QC HPLC Phenomenex Prodigy ODS column: 4.6 X 250 mm, 5 µm were purchased from Phenomenex. Sterile pyrogen-free filters were purchased from Millipore.

**Automated radiochemical synthesis of [$^{18}$F]DCP.** Radiochemical synthesis of [$^{18}$F]DCP was automated on the TRASIS AIO radiochemistry module, following the typical [$^{18}$F]Fluoride ([$^{18}$F]$^-$) nucleophilic substitution reaction. TRASIS AIO is an automated radiosynthesis module that produces GMP grade PET radiopharmaceuticals for human injections. The module includes two reactor vials that allow variable temperature control, internal radioactivity detector, temperature sensors, automated kit setup with syringes, vials and transparent loops. HPLC system includes one vacuum pump, one isocratic pump with a UV lamp. [$^{18}$F]$^-$ was produced from [$^{18}$O]$H_2O$ using a GE PETtrace 800 cyclotron through the $^{18}$O(p,n)$^{18}$F reaction at Wake Forest Cyclotron Facility. [$^{18}$F]$^-$ was directly trapped into a pre-conditioned QMA cartridge via a positive helium push. The residual radioactivity was then eluted from the cartridge using a reaction mixture solution, containing Kryptofix 222 (~7 mg) and $K_2CO_3$ (~2 mg) in a solution of anhydrous ACN:$H_2O$ (3:1) into the reaction vial. The residue was azeotropically dried at 110 °C under a nitrogen stream with anhydrous ACN (3 x 1.5 mL) to form [$^{18}$F]KF-K222 complex. After the reaction mixture was dried, the corresponding tosylate precursor (2.0 mg) in anhydrous DMF (350 µL) was added into the reaction vial and was heated at 110 °C for an additional 15 min. After cooling to room temperature, the reaction mixture was quenched with HPLC mobile phase (3.0 mL). The reaction mixture was then directly injected onto a reverse phase Phenomenex Luna C18 HPLC column (250 x 10 mm, 10 µm). The HPLC mobile phase was 52% ACN in 0.1 M aqueous ammonium formate buffer solution (pH 4.5) and elution was monitored at 254 nm with a flow rate of 5 mL/min. Under these conditions, [$^{18}$F]DCP was collected between 8-10 min directly into a vial containing sterile water (50 mL). The radioactive aqueous solution was then passed through a SepPak C18 Plus cartridge (Waters) to trap the [$^{18}$F]DCP. Further, the radioactive product was eluted with 10% absolute ethanol solution in saline. The final dose was filtered using a sterile 0.22 µm pyrogen-free Millipore filter into a sterile vial and used for cell culture, animal studies, and quality control analysis.

**Quality control analysis of [$^{18}$F]DCP.** The chemical and radiochemical purity of the collected radioactive aliquot was verified by performing a HPLC injection on a Phenomenex Prodigy ODS (4.6 X 250 mm, 5 µm). The mobile phase was 50% ACN: 50% 0.1 M ammonium formate buffer solution pH 7.0; the UV detection was set at 254 nm with a flow rate of 1.0 mL/min. Under these QC HPLC conditions, the single injection of [$^{18}$F]DCP showed retention time at 5.6 min. The radioactive peak was further authenticated by performing a co-injection with non-radioactive [$^{19}$F]DCP standard, which displayed similar retention times.

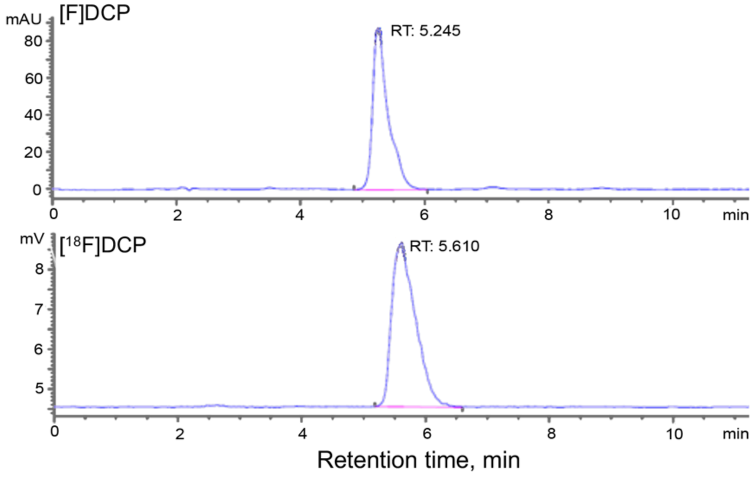


**Figure S4. Quality control analysis of [$^{18}$F]DCP showing consistent chromatographic properties with [$^{19}$F]DCP.**

**3. Chemical Synthesis of cold [$^{19}$F]DCP**

**3-ethoxy-6-(3-(1-(2-fluoroethyl)-*1H*-1,2,3-triazol-4-yl)propyl)cyclohex-2-en-1-one (3)**

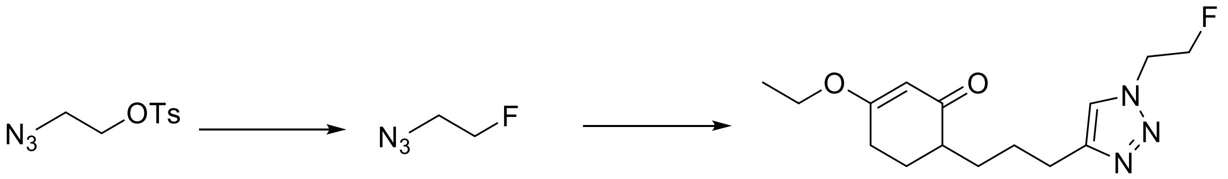


A solution of TBAF in THF (1 M, 3.0 mL, 3.0 mmol) was added to 2-azidoethyl-4-toluenesulfonate (0.245 g, 1.0 mmol) and the resulting solution stirred at room temperature overnight to yield 1-azido-2-

fluoroethane solution that was used in the next step without isolation or purification. 3-Ethoxy-6-(pent-4-yn-1-yl)cyclohex-2-en-1-one (0.105 g, 0.5 mmol), sodium ascorbate (24 mg, 0.12 mmol in 0.5 mL $H_2O$) and $CuSO_4$ (16 µl of 1 M solution in $H_2O$) were added to the solution of 1-azido-2-fluoroethane and the mixture was stirred overnight. TLC indicated the formation of a new product, the solvent was removed under vacuum to give a crude product that was purified by silica gel column chromatography (eluent: EtOAc) to give a yellow solid (0.102 g, 0.35 mmol. 70 % yield).

**Data:**

**$^1$H-NMR ($CDCl_3$) δ:** 7.39 (s, 1H), 5.23 (s, 1H), 4.79 (dd, $^1J$ = 6.0 Hz, $^2J$ = 3.0 Hz, 1H), 4.65 – 4.60 (m, 2H), 4.54 – 4.51 (m, 1H), 3.81 (dd, $^1J$ = 15.0 Hz, $^2J$ = 6.0 Hz, 2H), 2.68 (td, $^1J$ = 7.5 Hz, $^2J$ = 3.0 Hz, 2H), 2.34 (dd, $^1J$ = 7.5 Hz, $^2J$ = 4.5 Hz, 2H), 2.18 – 2.10 (m, 1H), 2.06 – 1.96 (m, 1H), 1.88 – 1.62 (m, 4H), 1.46 – 1.18 (m, 2H), 1.28 (t, $J$ = 7.5 Hz, 3H) (**Figure S5A**).

**$^{13}$C-NMR ($CDCl_3$) δ:** 201.42, 176.86, 148.36, 121.76, 102.17, 81.52 (d, $J$ = 171.0 Hz), 64.18, 50.36 (d, $J$ = 20.2 Hz), 44.94, 29.10, 28.07, 26.80, 26.27, 25.70, 14.15 (**Figure S5B**).

**LRMS-ESI$^+$ (m/z):** $[M + H]^+$ predicted 296.2, found 296.2.

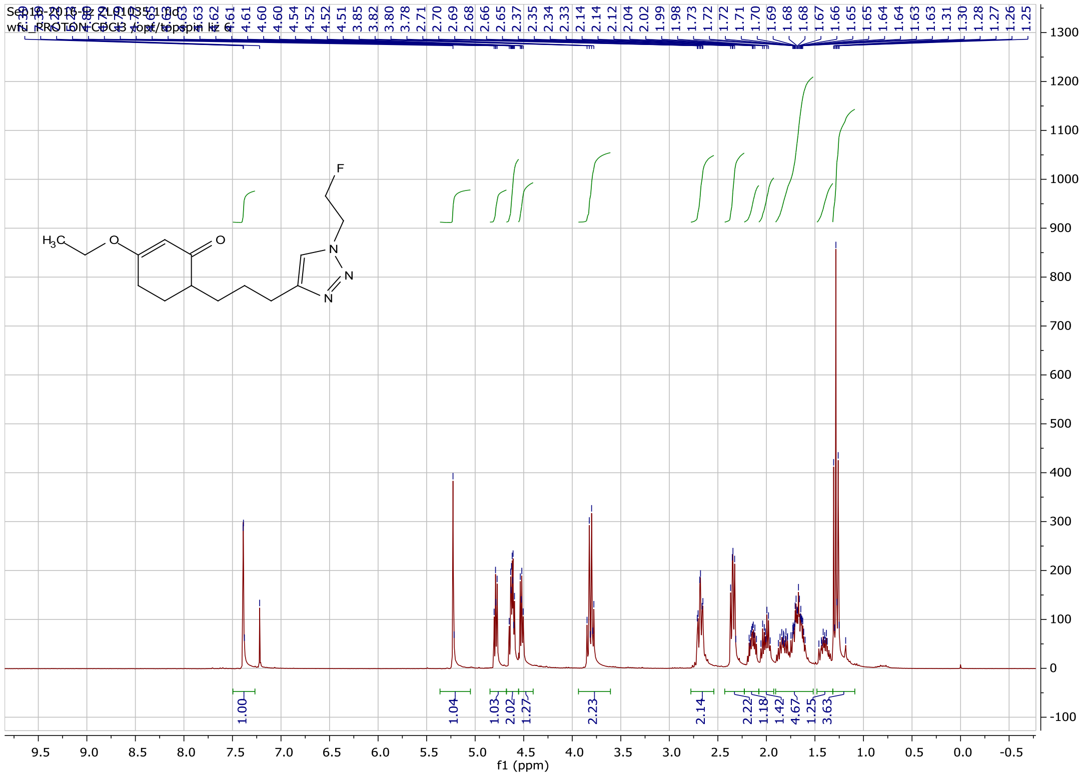


**Figure S5A. $^1$H-NMR spectrum of precursor 3 in [$^{19}$F]DCP synthesis.**

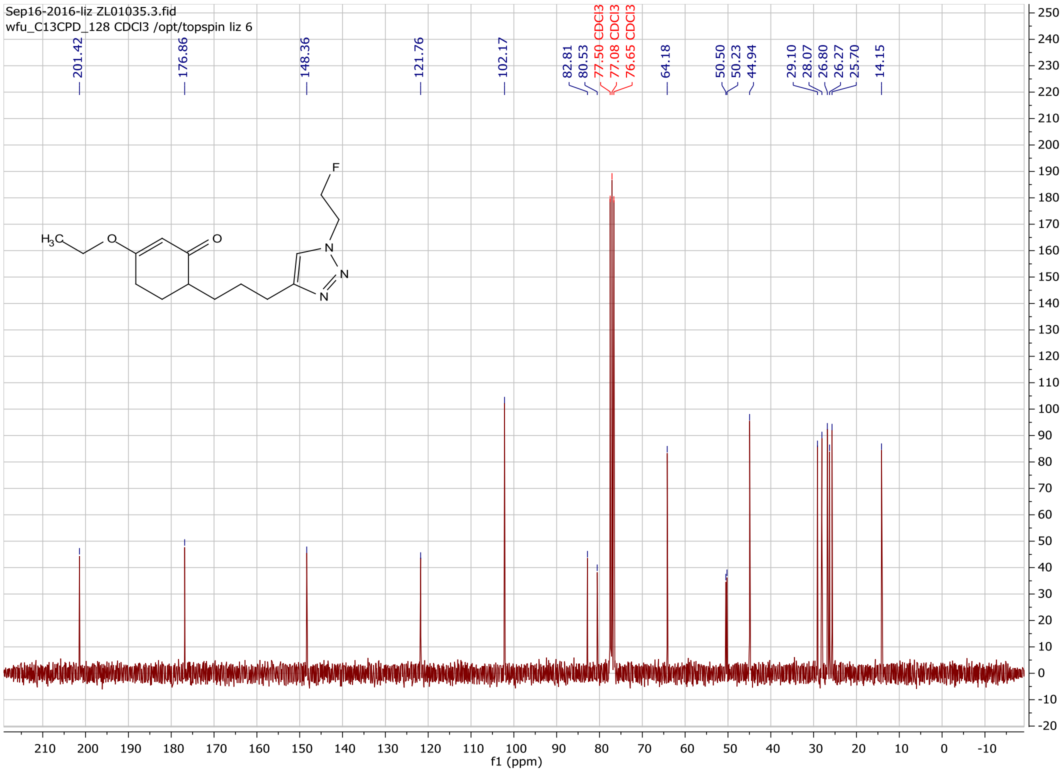


**Figure S5B. $^{13}$C-NMR spectrum of precursor 3 in [$^{19}$F]DCP synthesis.**

**4-(3-(1-(2-fluoroethyl)-*1H*-1,2,3-triazol-4-yl)propyl)cyclohexane-1,3-dione ([$^{19}$F]DCP)**

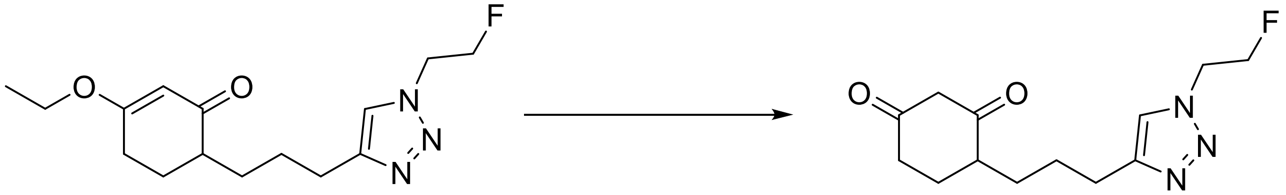

Ceric ammonium nitrate (CAN, 25 mg, 0.046 mmol) was added to a solution of 3-ethoxy-6-(3-(1-(2-fluoroethyl)-*1H*-1,2,3-triazol-4-yl)propyl)cyclohex-2-en-1-one (0.102 g, 0.35 mmol) in acetonitrile and water (3 mL : 3 mL) and the resulting solution was refluxed for 5 hours. Upon cooling, the solvent was removed under vacuum and the crude product was purified by C18 reversed phase silica gel column (elution: acetonitrile in water, 3% to 50%). Lyophilization of the appropriate fractions provided a light-yellow oil (0.066 g, 0.25 mmol, 71 % yield).

**Data: $^1$H-NMR ($CDCl_3$) δ:** 7.50 (d, $^1J$ = 6.0 Hz, 1H), 5.48 (s), 4.88 – 4.87 (m, 1H), 4.72 – 4.69 (m, 2H), 4.63 – 4.62 (m, 1H), 3.44 (d, $^1J$ = 6.0 Hz, 1H), 2.78 – 2.72 (m, 2H), 2.68 – 2.54 (m, 1H), 2.44 – 2.34 (m, 2H), 2.19 – 2.02 (m, 2H), 1.85 – 1.75 (m, 3H), 1.61 - 1.56 (m, 2H) (**Figure S6A**).

**$^{13}$C-NMR ($CDCl_3$) δ:** 204.63, 203.94, 194.34, 188.15, 148.02, 147.86, 122.16, 121.96, 81.55 (d, $J$ = 171.0 Hz), 58.33, 50.55 (dd, $^{1}J$ = 32 Hz, $^{2}J$ = 4.5 Hz), 41.47, 39.67, 30.24, 29.62, 28.58, 26.78, 26.66, 26.06, 25.46, 25.36, 24.47 (**Figure S6B**).

**$^{19}$F-NMR ($CDCl_3$) δ:** -221.56, -221.59 (**Figure S6C**).

**LRMS-ESI$^+$ (m/z):** $[M + H]^+$ predicted 268.1, found 268.0.

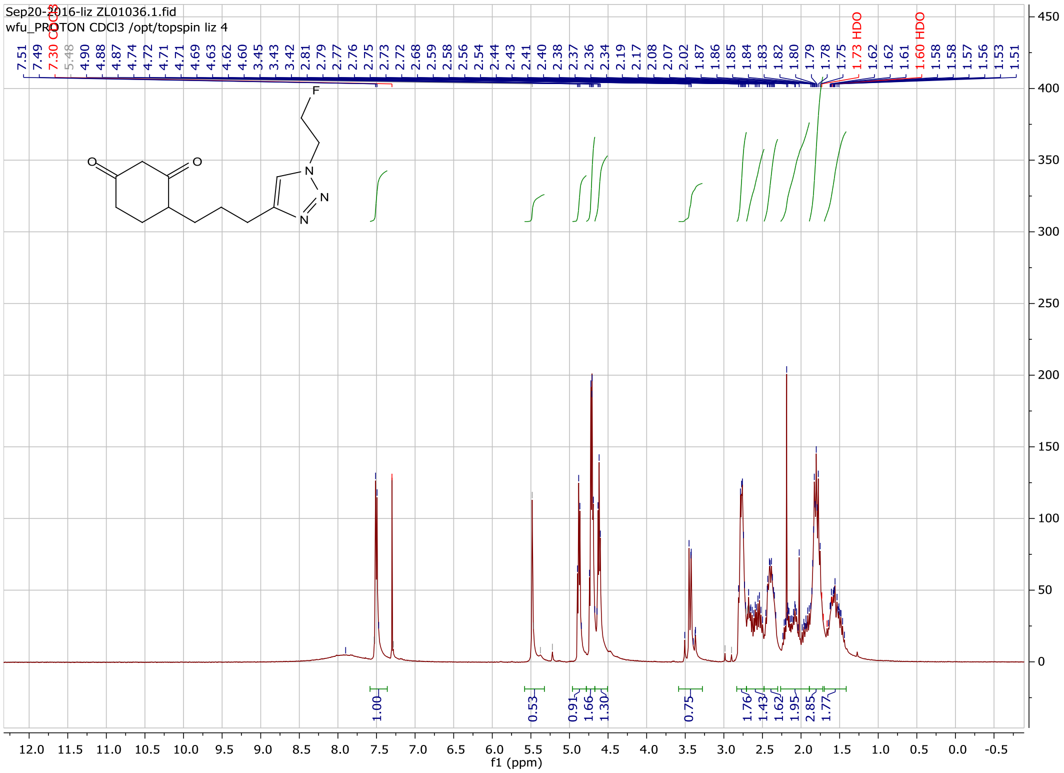


**Figure S6A. $^{1}$H-NMR spectrum of [$^{19}$F]DCP.**

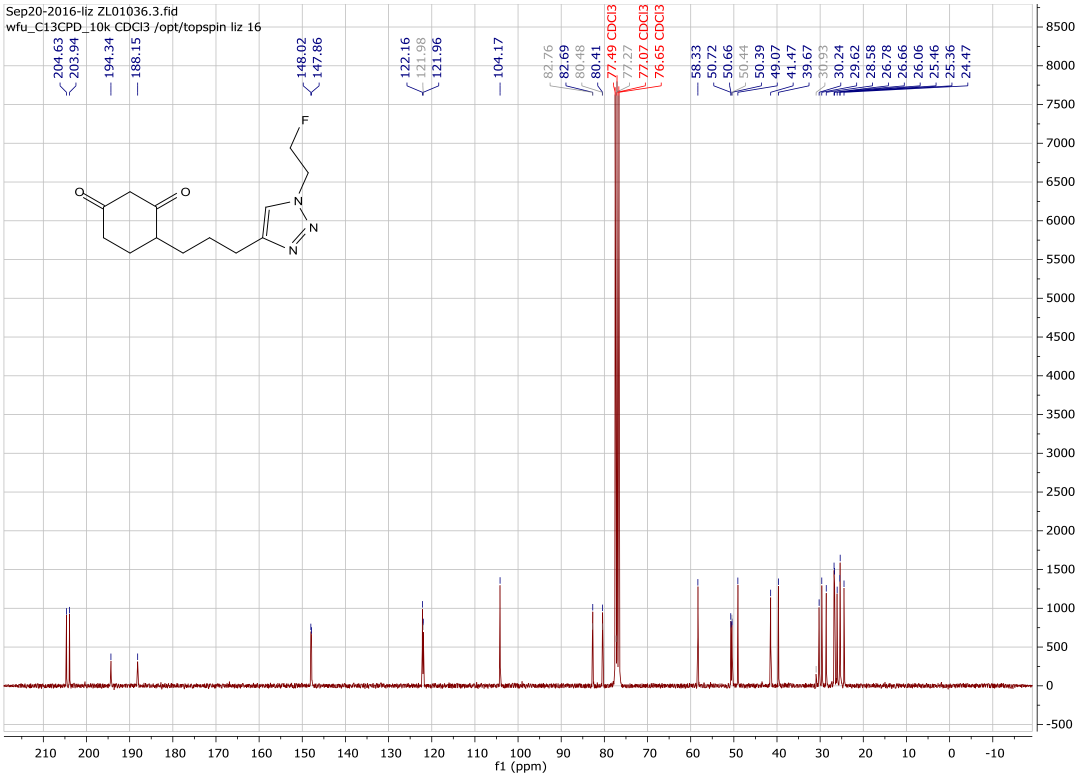


**Figure S6B.** $^{13}$**C-NMR spectrum of [**$^{19}$**F]DCP.**

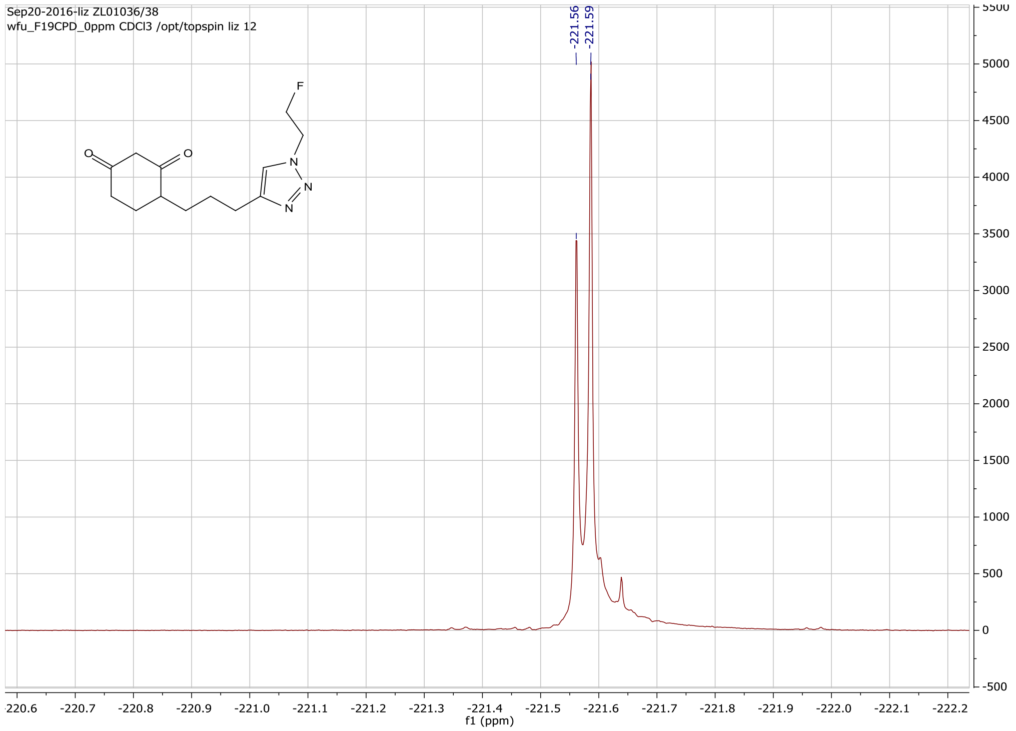


**Figure S6C.** $^{19}$**F-NMR spectrum of [**$^{19}$**F]DCP.**
